# Decoupling individual host response and immune cell engager cytotoxic potency

**DOI:** 10.1101/2024.06.22.600188

**Authors:** Cristina Gonzalez Gutierrez, Adrien Aimard, Martine Biarnes-Pélicot, Brigitte Kerfelec, Pierre-Henri Puech, Philippe Robert, Francesco Piazza, Patrick Chames, Laurent Limozin

## Abstract

Immune cell engagers are molecular agents, usually antibody-based constructs, engineered to recruit immune cells against cancer cells and kill them. They represent a versatile and powerful tool for cancer immunotherapy. Despite the multiplication of new engagers tested and accepted in the clinics, how molecular and cellular parameters influence their action is poorly understood. In particular, disentangling the respective role of host immune cells and engager biophysical characteristics is needed to improve their design and efficiency. Focusing here on harnessing antibody dependent Natural Killer cell cytotoxicity, we measure the efficiency of 6 original bispecific antibodies (bsAb), associating an anti-HER2 nanobody and an anti-CD16 nanobody. *In vitro* cytotoxicity data using primary human NK cells on different target cell lines exposing different antigen densities were collected, exhibiting a wide range of bsAb dose response. In order to rationalize our observations, we introduce a simple multiscale model, postulating that the density of bsAb bridging the two cells is the main parameter triggering the cytotoxic response. We introduce two new microscopic parameters: the surface cooperativity describing bsAb affinity at the bridging step and the threshold of bridge density determining the donor-dependent response. Both parameters permit to rank Abs and donors and to predict bsAb potency as a function of antibodies bulk affinities and receptor surface densities on cells. Our approach thus provides a general way to decouple donor response from immune engagers characteristics, rationalizing the landscape of molecule design.

---

Antibodies are powerful therapeutics used in a wide range of diseases^1^ and have revolutionized cancer treatment by harnessing the immune system against tumour cells.^2^ However, patients do not respond equally to immunotherapies^3^ and deconvoluting the contribution of therapeutic molecules and patient cells to the response is needed to improve Abs design and efficiency. Also, conventional Abs mode of action is complex since those divalent molecules can bind one or two antigens *via* their Fab fragments, as well as immune cell receptors *via* their Fc fragment. As an example, trastuzumab, the reference therapeutic antibody used against HER2 positive breast cancer,^4^ can act simultaneously by blocking antigen receptors, and by recruiting immune cells displaying Fc receptors, like CD16.^5^

Antibodies can be engineered as bispecific molecules targeting directly two different epitopes or antigens.^6^ Immune cell engagers are engineered antibodies redirecting immune cells, entailing ease-of-use and multifunctional arms.^7^ Motivated by applications in human health, a lot of new molecules are continuously produced and evaluated.^8–10^ They can target different immune cells such as T or NK cells, or target multiple tumour antigens for improved specificity.^7^ T cell engagers are among the most developed and potent molecules,^11, 12^ and may entail important side effects, like cytokine release syndrome.^13^ A way to diminish unwanted side cytotoxicity is by reducing the affinity to TCR/CD3,^14^ but the related mechanisms are poorly known. Some other engagers target NK cells,^15–18^ often *via* the activating Fc receptor CD16^19, 20^ and triggering antibody-dependent cytotoxicity (ADCC), and are less prone to excess of cytokine production.^21^

Despite their potential, these molecules are laborious to test and characterize, and their efficiency may vary between patients, highlighting the need for establishing and rationalizing design principles. Avidity effects play a central role in Ab binding and efficiency,^22^ but are difficult to predict.^23, 24^ Some models have been proposed to quantify avidity in a *cis* configuration,^25, 26^ and *trans* configuration was examined numerically for model liposomes.^27^ Additionally, multi-scale models of cytotoxicity mediated by Abs^28–31^ are based on partial differential equations or on numerical simulations, but without estimating microscopic parameters based on systematic measurements.

Ultimately, engagers are working by enforcing the formation of an immune synapse between the immune cell and the tumour cell.^32, 33^ In this intercellular cleft of typically 10-30 nm thickness, the roles of affinity, valency, geometry and force are interwined to trigger immune signaling,^34^ controlling the cytotoxic function. While the natural immune synapse for T cells or NK cells is well studied, the microscopic parameters triggering synapse establishment are still debated.^35, 36^ Additionally, synapses established by T cell engagers might differ from natural ones and are poorly characterized.^37^ Thus, engagers could also be harnessed to probe the biology and biophysics of the synapse.^34^

Overall, the quantitative basis evaluating binding and efficiency of engagers is largely missing, leaving fundamental questions opened: what is the concentration of engager required to get half efficacy, i.e. engager potency? How does potency depend on effector and target cells, as well as antibody binding properties? In this study, we designed a defined set of six original bispecific NK engagers (bsAb) associating two nanobodies using a fixed format.^38, 39^ Based on systematic measurements of their ADCC potency using 15 different donors and 3 different target cells, we introduce a multiscale model taking into account microscopic binding parameters at the synapse and the density of bridging bsAb that sets the effector cell response. This approach reveals the dependence of bsAb potency as a function of the affinity of each Nanobody (Nb, or single domain antibody) forming the bsAb and the receptor density on cell partners. Emerging from the model, two new parameters define bsAb cooperative binding and donor-dependent NK cell sensitivity. They permit to decouple molecular and cellular determinants of cytotoxic response.

## Results

### Picomolar bsAb potency (EC_50_) depends on both antibody and donor

The lysis of tumour cells by NK cells was investigated in a co-culture of adherent target cells expressing variable HER2 densities (using 3 different cell lines: MCF7-WT, SKBR3 and MCF7-HER2+, see Tab. S1) and primary human NK cells isolated from 15 healthy donors (noted A-O in chronological order of the experiments) (Fig. 1A). We tested, in each case, 6 bispecific antibodies based on the previously described bsFab (for Fab-like bispecific) format,^40^ constructed by association of one anti-HER2 nanobody (among clones CE4, CA5 and C7b) and one anti-CD16 nanobody (among clones C21 and C28^38, 41, 42^) fused using the human C*κ*/CH1 heterodimerization motif (see measured affinity parameters in Tab. S2). The fraction of target cells lysed after an overnight incubation with primary NK cells was measured as a function of the concentration of bsAb (*c*) in the culture using a luminescent cell viability assay.^38^ Following characterization standards for monoclonal antibodies, we fit each individual lysis curve via a Hill function (setting the usual Hill exponent to 1):

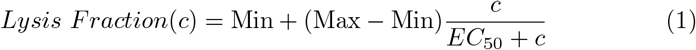

where Min, Max are the minimum and maximum values of the total lysis fraction, respectively, and EC_50_ is the bsAb potency, illustrated on an example of measurement on Fig. 1B. All lysis data and superimposed fitting curves are shown in Fig. S1.

**Figure 1:**
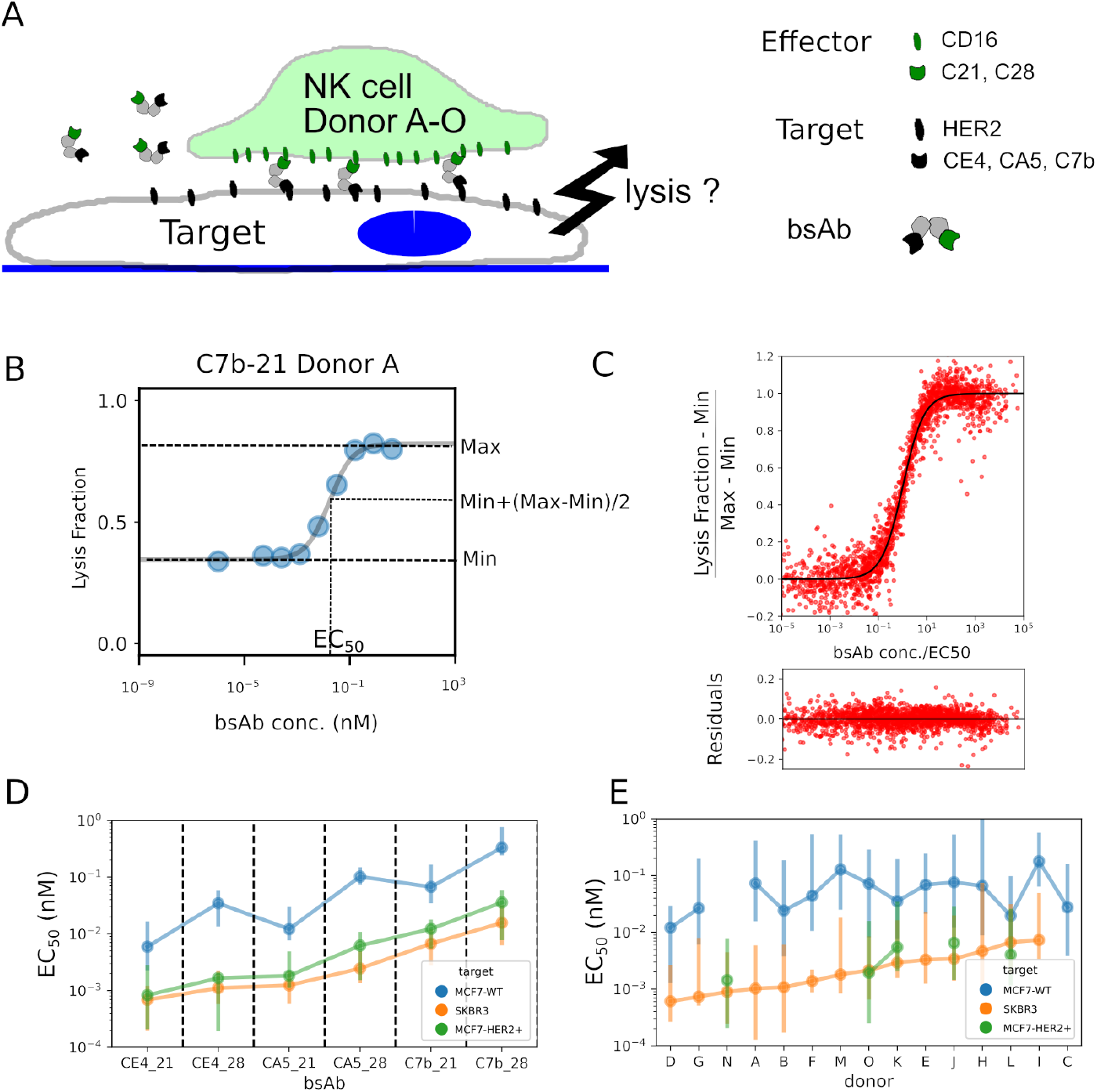
ADCC assay and dependence of potency (EC_50_) on bispecific antibody (bsAb) and NK cell donor. A. Schematic representation of co-culture assay mixing adherent tumour target cells (MCF7-WT, SKBR3 or MCF7-HER2+) and primary human NK cells from 15 donors in the presence of bsAb. Six bsAb anti-HER2*×* CD16 constructs are tested, based on 3 anti-HER2 (Nanobodies CE4, CA5 or C7b) and 2 anti-CD16 (Nanobodies C21 or C28). B. Example of lysis fraction *vs* bsAb concentration measured on the MCF7-HER2+ target cell line and bispecific Ab C7b-21, donor *A*. Data was fitted with Eq. (1) (black line). C. Result of Hill fit for all conditions. Data are normalized using the fitting parameters Min, Max and EC_50_ and compared to the normalized Hill function *c/*(1 + *c*) with c in nM units (black line). Raw residuals are shown below. Each dot represents one Donor/bsAb/target condition (198 conditions). D. EC_50_ for each target cell line and bsAb. Each point is the median on the donors, with the bar representing 95% percentile interval. bsAbs are ranked according to the median value for SKBR3 cell line. E. EC_50_ for each target cell line and donor. Each point is the median on the bsAbs, with the bar representing 95% percentile interval. Donors are ranked according to the median value for SKBR3 cell line.

The goodness of the fits can be assessed visually on Fig. 1C, after normalizing the lysis curves, vertically with respect to the minimal and maximal lysis fraction and horizontally with respect to EC_50_. The fit residuals shown under the curve were homogeneously distributed with a standard deviation of 0.041. The bestfit values of the free parameters are reported for all conditions in Fig. 1D-E for EC_50_ and Fig. S2 for Min and Max. Min and Max appear to be negligibly impacted by the nature of the bsAb and to be correlated for each target cell line (Fig. S3). Remarkably, all values of bsAb potency were found to be in the picomolar range (Figs. 1D-E), 2 or 3 orders of magnitude smaller than the affinities that quantify the strength of binding to their respective epitopes (Tab. S2). As expected, the EC_50_ obtained for the cell lines SKBR3 and MCF7-HER2+ were much smaller than those found for MCF7-WT, correlating with the expression level of HER2. For each target cell line, the median was calculated either on donors and displayed by bsAb (Fig. 1D) or calculated on bsAbs and displayed by donor (Fig. 1E). bsAb and donors were ranked by the median EC_50_ obtained on SKBR3. The robustness of ranking with EC_50_ was quantified using multiple Spearman rank-tests, as described in the Supplementary Material section. EC_50_ provides a strong ordering of both Abs and donors; Min and Max provide a strong ordering of donor only (Fig. S4 and Table S3 upper half). We also systematically compared bsAb or donors pairwise using non-parametric Wilcoxon test (Fig. S5). Differences are significant between most bsAbs, and only between about half of the donors. When comparing parameters for the same donor and bsAb on different cell lines, we find a high correlation for EC_50_ (Fig. S6). Taken together, those results suggest that it may be possible to decouple the influence of bsAb and donor on the potency.

### Multiscale model of bispecific dependent cell mediated cytotoxicity

In order to provide a microscopic interpretation of the Hill parameters Min, Max and EC_50_, we propose to describe the role of bispecific Abs on NK cytotoxicity by integrating 3 different scales: i) at the molecular scale, the bispecific antibodies bind to the HER2 (tumour cell side) and the CD16 (effector cell side) antigens at the immune synapse; ii) at the cellular scale, the rate of ADCC depends on the density of bridging bsAb; and iii) at the sample scale, the amount of target cells surviving over time is set by the (overall) killing rate, which is the sum of the spontaneous killing and antibody-dependent killing.

At the molecular scale, the bsAb binding to the membrane receptors occurs in two steps, from solution to cell surface (step A) and from cell surface to cell surface (step B) (Fig. 2A). The dissociation constants *K*_1_ and *K*_2_ for the first step A have the dimension of a concentration, whereas the dissociation constants *K*_*S*1_ and *K*_*S*2_ for the second step B have the dimension of a surface density (Tab. 1). Separating the molecular and cellular time scales,^28^ we suppose that the reactions are at equilibrium, and occur in the synaptic cleft with a fixed inter-membrane distance *h* = 10 nm. We introduce the cooperativity parameter *α* which relates volume and surface dissociation constants *via K*_*Si*_ = *hK*_*i*_*/α*; it represents the gain (*α* ≫ 1) or loss (*α* ≪ 1) of affinity for R_2_ when binding occurs after attachment to target R_1_ (step B), compared with the affinity for R_2_ when binding occurs first, directly from the solution (step A). A similar cooperativity parameter was previously proposed to describe tripartite equilibria^43^ or divalent ligand-monovalent molecule binding.^44^

**Figure 2:**
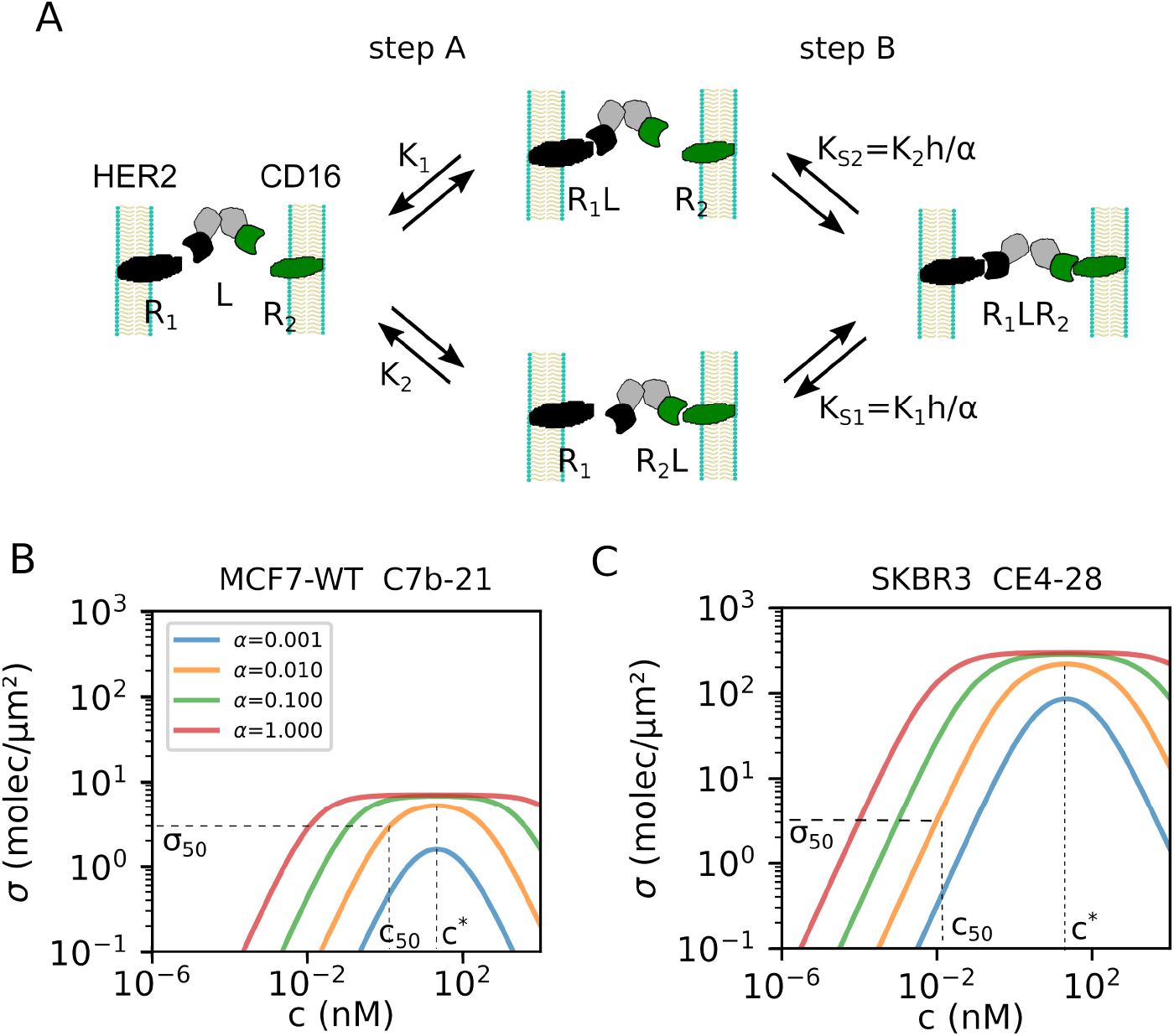
Physical model for bsAb dependent cell-mediated cytotoxicity. A. Reaction scheme for the two-step binding of bispecific antibody *L* at the immune synapse on membrane receptors *R*_1_ (tumour side, HER2) and *R*_2_ (effector side, CD16). *K*_1_, *K*_2_, *K*_*S*1_, *K*_*S*2_ are dissociation constants. B-C. The equilibrium density *σ* of bridging bsAbs as a function of bulk bsAb concentration *c* given by the solution of Eq. (S1). Each colored curve has been obtained for a different value of the cooperativity parameter *α* varied from 0.001 to 1. Other parameters *K*_1_, *K*_2_, Σ_1_, Σ_2_ correspond to Target MCF7-WT and antibody C7b-21 (B) or Target SKBR3 and antibody CE4-28 (C) (values in Tabs. S1 and S2). The horizontal line in (B,C) represents an example of *σ* threshold value for the NK cell cytotoxic response (here *σ*_50_ = 3 molec/*µ*m^2^). Together with the value of the cooperativity (here *α* = 0.01), it sets the value of *c*_50_, such that *σ*(*c*_50_) = *σ*_50_. The maximum density *σ*^∗^ is found for 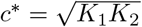.

The equilibrium density *σ* of bridging bsAb at the synapse is given by the solution of Eq. S4 derived using Tab. S4. The computed *σ*(*c*) is illustrated for bispecific Abs C7b-21 on MCF7-WT target cells in Fig. 2B and for bispecific Abs CE4-28 on SKBR3 target cells in Fig. 2C. Its maximal value *σ*^∗^ is obtained at the concentration 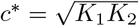 and is bounded from above by the minimum of receptor densities on each side Σ_min_=Min(Σ_1_,Σ_2_). The figure S7 illustrates the dependence of *σ*(c) on different parameters of the model. It shows the classical hook effect for *c* ≥ *c*^∗^,^43, 45^ as well as how the maximal bridging density is set by Σ_min_ for large values of *α*, but reduces with *α*.

At the cell scale, the lysis is initiated by complex binding processes involving many receptors on both sides of the immune synapse. We approximate the overall lysis rate to be the sum of the rate of spontaneous lysis *k*_0_, independent of the presence of antibodies, plus a second term that describes the bsAbs-dependent cell cytotoxicity. In the simplest scenario, one may postulate that this term takes the simple Michaelis-Menten form, with maximal rate *k*_*L*_ and half-rate obtained at density *σ*_50_ of bridging bsAb. Overall, we have:

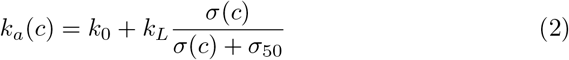

The titration of the overall lysis rate takes its maximum value at *σ*^∗^ = *σ*(*c*^∗^), so long as the bsAb concentration range includes 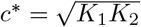 The concentration *c*_50_, such that *σ*(*c*_50_) = *σ*_50_, is indicated on Fig. 2B,C and will be considered later in the study. Comparison of the cases of MCF7-WT (Fig. 2B) and SKBR3 (Fig. 2C) shows that the same surface density of bridging bsAb *σ*_50_ is reached at a much lower concentration *c*_50_ on a high antigen density target.

At the cell population scale, the variation with time of the number of living target cells, *T* (*t*), depends on the number of effector cells *N*_0_, the total lysis rate *k*_*a*_ and on the fraction of active effector cells, which we term *γ*(*t*). Note that target cells may still proliferate with a rate *k*_*p*_. The lysis fraction as a function of time Eq. S6 is derived in the Suppl. Mat. Taking an exponential decay for the function *γ*, this equation fits very well with the lysis data obtained by Real Time Cytoxicity Assay (RTCA, xCelligence, Agilent) based on the follow up of impedence generated by adherent target cells (Fig. S8A). The dependence of the lysis rate as a function of bsAb concentration exhibits a saturation (Fig. S8B), which justifies *a posteriori* the form taken in Eq. 2, as discussed later.

### Molecular and cellular parameters from fitting of lysis data

Our dataset comprises ADCC response curves measured for 3 target cell lines, each comprising NK cells from up to 15 donors, for the 6 bsAb constructs considered. This gives a total of 33*×*6=198 biological conditions. For each target cell line separately, we performed a global fit with 4 parameters for each *lysis fraction* vs *bsAb concentration* set (see Fig. S1). The data are directly fitted without normalization using Eqs. (2) and Eq. (S6). Parameters *k*_0_ and *k*_*L*_ represent the spontaneous lysis rate and the maximal bsAb-dependent lysis rate, respectively, according to Eq. (2). The two other free parameters are the surface density threshold of bridging bsAbs, *σ*_50_, and the cooperativity for the bridging, *α*. In order to reduce the total number of parameters, we made two crucial hypotheses: (i) The cooperativity parameter *α* is a specific signature of the molecular architecture of each type of bispecific antibody; and (ii) The lysis rates *k*_0_ and *k*_*L*_, and the threshold for bsAb-activated lysis, *σ*_50_, depend on the specific molecular details of the surfaces of NK and target cells. As a consequence, these three parameters depend on the donor but not on the bsAb. Supporting the two hypotheses above, a model-free analysis of the data (see Suppl. Mat and Fig. S9) shows that EC_50_ can be retrieved as the product of one bsAb dependent term and one donor dependent term. In principle, the parameter *k*_*L*_ may bear a signature of the specific antibody. However, we have verified that allowing the parameter *k*_*L*_ to depend on the bsAb does not impact significantly on the results.

The goodness of the fits can be assessed visually by looking at Fig. S10A, after normalizing as previously (Fig. 1B). A linearized version of the model (valid for low bsAb concentration) provides very similar results (Fig. S10B). This limit will be exploited in the next section. In both cases the residuals are homogeneously distributed and the standard deviation is 0.063, a value only marginally higher than the one obtained with independent Hill fits.

All best-fit values for parameters *σ*_50_ and *α* are reported in Fig. 3. As a first important observation, we note that cell-response density thresholds *σ*_50_ lie in the range 0.1-10 molecules/*µ*m^2^ for all donors, with values significantly lower for MCF7-WT (Fig. S11B). The values of bsAb cooperativity *α* lie typically between 0.01 and 0.1, denoting a strong hindrance for the second binding step at the synapse. We note that *α* is not siginificantly varying with the target cell line (Fig. S11A). While *α* appears weakly dependent on the bsAb on MCF7-WT, values are strongly correlated between the cell lines exhibiting high HER2 density (Fig. S12C). The best-fit values of *k*_0_ and *k*_*L*_ are reported in Fig. S13. It can be noticed that values of *k*_0_ are comprised between 0.1 and 0.75 (in units of *N* (0)*A*(*T*)), spanning a comparable range for the 3 target cell lines, while *k*_*L*_ values vary typically between 0.4 and 4 (in units of *N* (0)*A*(*T*)). This suggests that the maximum relative increase of the killing rate due to the bsAb-induced bridging, *k*_*L*_*/k*_0_, is near 400 % on average, which underlines the interest of NK cell engagers for therapy.

**Figure 3:**
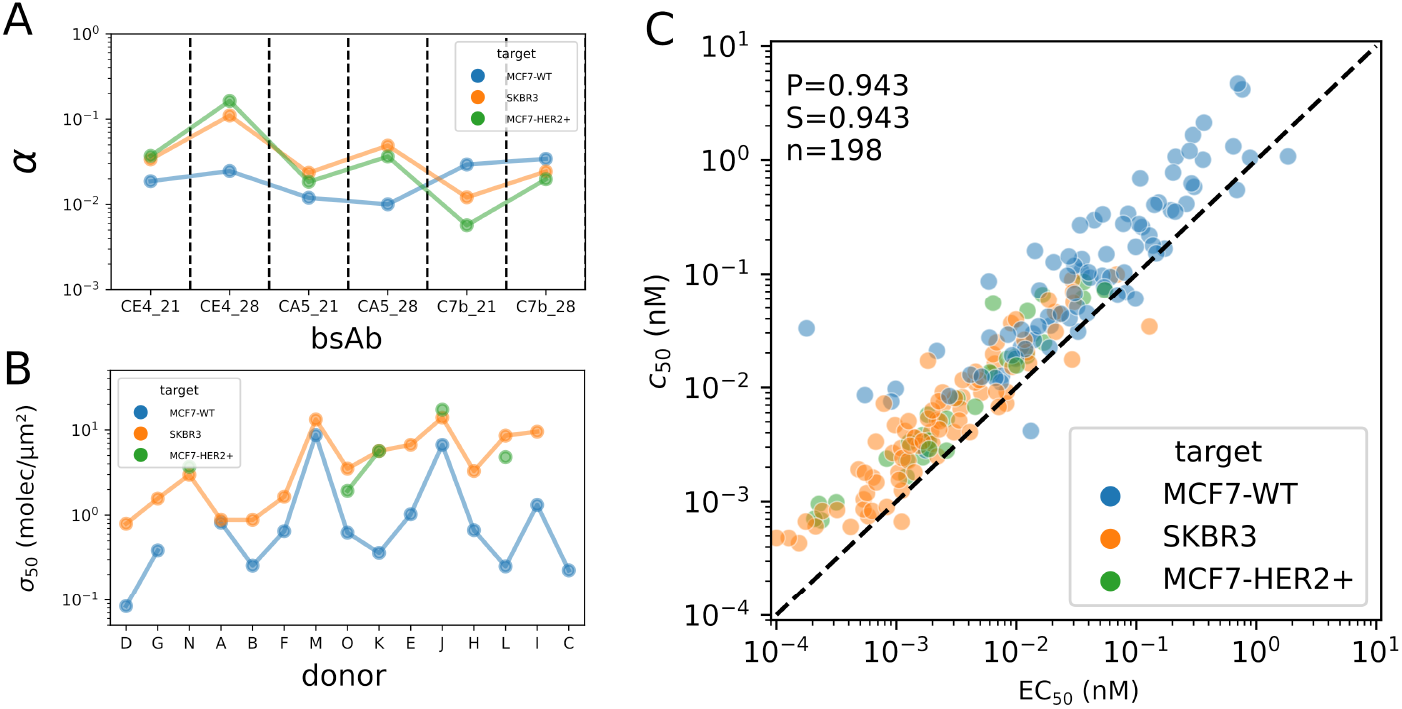
bsAb cooperativity *α*, NK cells threshold response *σ*_50_ and estimator of potency *c*_50_. A. Best-fit values *α* for each bsAb. B. Best-fit values *σ*_50_ for each donor. C. Relation between values of EC_50_ measured *via* the Hill analysis (horizontal axis) and the potency estimator *c*_50_ from Eq. 3 (vertical axis). Each point represents one Donor/bsAb/Target condition. Dashed lines represent *y* = *x*. P: Pearson coefficient. S: Spearman coefficient. n: number of points.

**Table 1:**
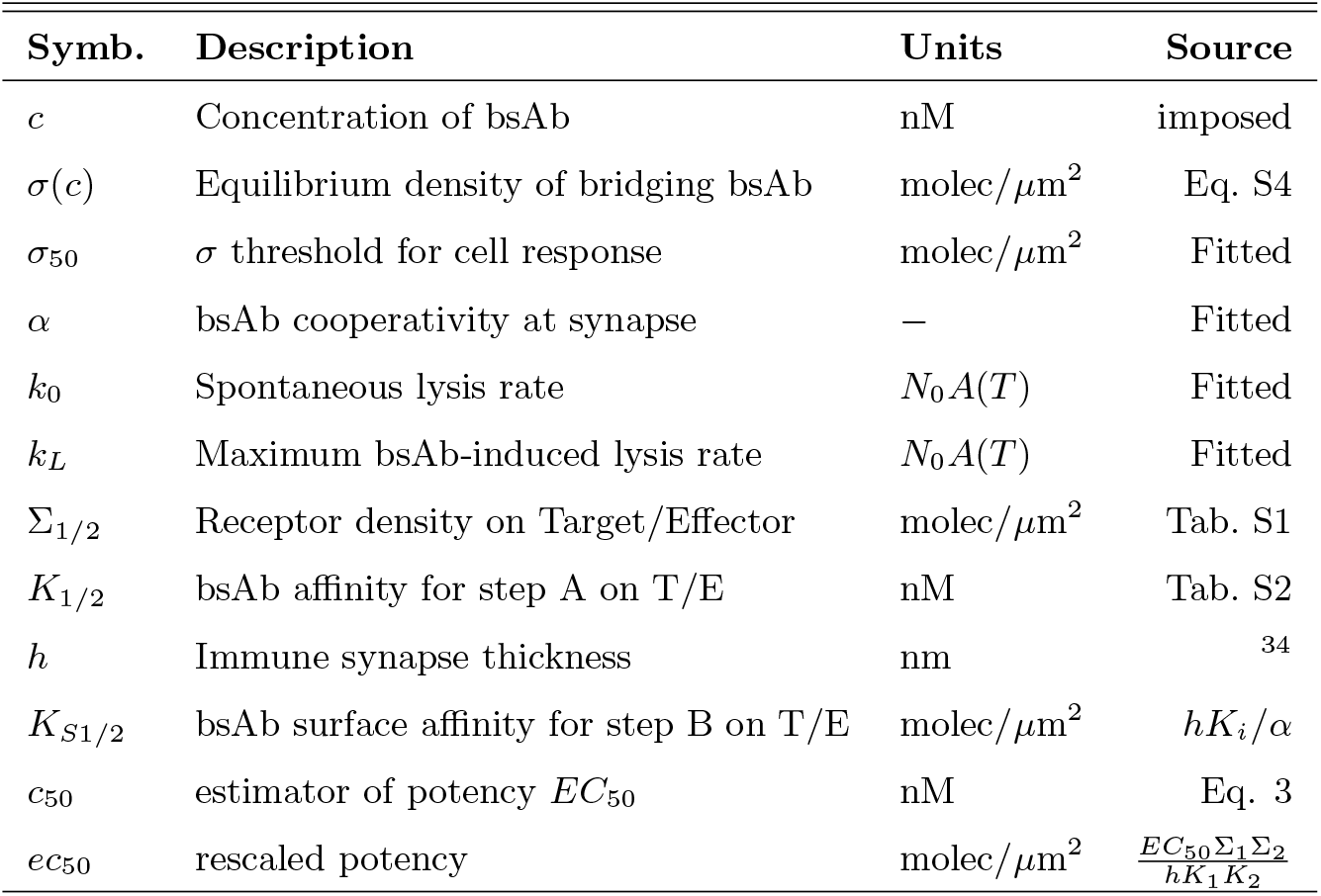
Variables used for the multiscale cytotoxicity model. T: Target. E: Effector. Steps A and B refer to the reaction scheme of Fig. 2A.

| Symb. | Description | Units | Source |
| --- | --- | --- | --- |
| $c$ | Concentration of bsAb | nM | imposed |
| $\sigma(c)$ | Equilibrium density of bridging bsAb | molec/$\mu\text{m}^2$ | Eq. S4 |
| $\sigma_{50}$ | $\sigma$ threshold for cell response | molec/$\mu\text{m}^2$ | Fitted |
| $\alpha$ | bsAb cooperativity at synapse | — | Fitted |
| $k_0$ | Spontaneous lysis rate | $N_0 A(T)$ | Fitted |
| $k_L$ | Maximum bsAb-induced lysis rate | $N_0 A(T)$ | Fitted |
| $\Sigma_{1/2}$ | Receptor density on Target/Effector | molec/$\mu\text{m}^2$ | Tab. S1 |
| $K_{1/2}$ | bsAb affinity for step A on T/E | nM | Tab. S2 |
| $h$ | Immune synapse thickness | nm | <sup>34</sup> |
| $K_{S1/2}$ | bsAb surface affinity for step B on T/E | molec/$\mu\text{m}^2$ | $hK_i/\alpha$ |
| $c_{50}$ | estimator of potency $EC_{50}$ | nM | Eq. 3 |
| $ec_{50}$ | rescaled potency | molec/$\mu\text{m}^2$ | $\frac{EC_{50}\Sigma_1\Sigma_2}{hK_1K_2}$ |

**Table 2:** Prediction of potency compared with literature data. EC_50_ or EC_50_ ratios where predicted using Eq. 4 and compared to published data.

| # | Format | Ab / Ref. | $K_1$<br>(nM) | $K_2$<br>(nM) | Predicted<br>$EC_{50}$ (nM) | Measured<br>$EC_{50}$ (nM) |
| --- | --- | --- | --- | --- | --- | --- |
| 1 | Mab | Trast.<sup>38</sup> | 0.3 | 200 | 0.02 | $\sim 0.03$ |
| 2 | bsFab | CA5-21 * | 10 | 3 | 0.01 | 0.01 |
| | | | Param. | Ratio | Predicted<br>$EC_{50}$ ratio | Measured<br>$EC_{50}$ ratio |
| 3 | K-H | HER2×CD3<sup>53</sup> | $\Sigma_1$ | 100 | 0.01 | $\sim 0.01$ |
| 4 | K-H | HER2×CD3<sup>53</sup> | $K_1$ | 76 | 76 | $\sim 50$ |
| 5 | K-H | HER2×CD3<sup>53</sup> | $K_1$ | 47 | 47 | $\sim 50$ |

Trast.: Trastuzumab. K-H: Knob-Hole. \*: this study

### Scaling of potency at low bsAb concentration

The analysis of our dataset performed with the Hill function revealed that the bsAb potency was systematically below the bulk affinities that characterize single bonds on either side of the synapse, that is, EC_50_ ≪ *K*_1_, *K*_2_.

This observation suggests to consider the solution of Eq. S4 in the limit where 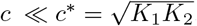.In this regime, the density of bridging bsAb takes the simple form: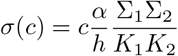, so that Eq. 2 becomes : 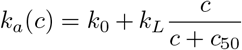,where *c*_50_ was previously introduced as the concentration of bsAbs that corresponds to the cell threshold density *σ*_50_, *ie σ*(*c*_50_) = *σ*_50_. At low bsAb concentration, we therefore obtain:

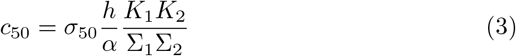

Note that the dependence of the lysis rate *k*_*a*_(*c*) on concentration is consistent with the RTCA measurements (Fig. S8B), justifying *a posteriori* the form of Eq. 2. Fig. 3C shows that *c*_50_ predicts satisfactorily the measured potency EC_50_ obtained through the Hill analysis, over more than 4 orders of magnitude. One notices however that predicted values are slighty higher than the measured ones. A more accurate prediction of EC_50_ can be obtained with a modified expression for *c*_50_ that also takes into account the value of *k*_*L*_ (see Fig. S14). While *c*_50_ is a good estimator of EC_50_, it requires that we know the values of *σ*_50_ and *α*. This can be partly addressed by considering the rescaled potency 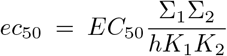, homogeneous to a surface density, which simplifies to *ec*_50_ ≃ *σ*_50_*/α* by using *c*_50_ as the estimate of *EC*_50_. The median *ec*_50_ per donor (respectively per bsAb) provides a relative estimate of *σ*_50_ (respectively *α*) (Fig. S15).

We can now decompose the estimate of the potency (Eq. 3) as a product of three terms that describe the separate contribution of the different players on ADCC:

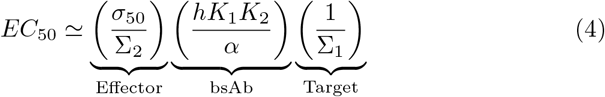

Their values are plotted on Fig. S16. The term for bsAb displays consistent values between the 3 target cell lines and overall a clear ranking of bsAb. The term for effector is very similar between targets with high antigen densities, and in average smaller on MCF7-WT, suggesting a higher specific contribution of effector to the potency on this target cell line. The specific impact of each nanobody constituting the bsFab, either on target or effector side, is shown on Fig. S17. Interestingly, on the effector side, the cooperativity contribution, *α*, of C28 is higher than that of C21, in line with the catch bond behavior identified previously for C28-CD16 bond.^42^

## Discussion

A better knowledge and understanding of the parameters influencing the antibody-mediated mechanism of ADCC would help rationalize the design of new therapeutic molecules. In this work, we have synthetized a set of six new nanobody-based NK cell engagers using a unique format (the 50 kDa bsFab format using the human IgG1 CH1 and Ck domain as heterodimerization motif), three nanobodies against the classical tumour associated antigen HER2 and two nanobodies targeting the activating receptor CD16. We systematically measured their potency in triggering the cytotoxicity of resting NK cells from human healthy donors on tumor cells expressing various levels of HER2. Our study provides a scaling of the potency as a function of measurable parameters such as affinities and receptor densities, revealing their combined roles. Two new parameters characterizing the bispecific bridging Ab on the one hand (*α*) and NK cell response on the other hand (*σ*_50_) were extracted, a first step towards decoupling molecular (e.g. drug-related) and cellular (e.g. patient-related) factors impacting cancer therapies based on immune cell engagers.

At the molecular scale, the bsAb cooperativity factor, *α*, was found to be about 0.01, which indicates a strong reduction of bond affinity for the bridging step at the synapse, as compared to the bulk affinity, which could be explained either by a lower on-rate or a higher off-rate. The on-rate can be influenced by the distance between the two paratopes of the bsAb, the epitope locations and accessibility on the antigens, which could become critical in order to match the binding length with the size of the synaptic cleft^46^ or by the restricted receptor diffusion within the membrane. The off-rate could be affected due to pulling forces exerted by target and effector cells on each other.^42, 47^ A characteristic value of *α* for each bsAb impacts cytoxicity at high HER2 density, but can not be precisely inferred at low HER2 density (Fig. 3A, S12C). This suggests that the decoupling between molecular and cellular parameters is more efficient at high HER2 density, possibly because the ADCC effect becomes then dominant compared to natural cytotoxicity.

At the cell scale, the effector threshold response *σ*_50_ was found to vary between 0.1 and 10 molecules/*µ*m^2^, much lower than the receptor densities on the cells (Tab. S2). Thus, on the effector side, the ratio with the CD16 receptor density, *σ*_50_*/*Σ_2_, lies between 0.1 and 1%, making bsAb a sensitive trigger of NK cell response. Integrated on the surface area of the synapse (typically 100 *µ*m^2 35^), it represents between 10 and 1000 bridging molecules. These estimates could be modulated by the receptor mobility within the plasma membrane,^48^ while we assume uniform densities in our model. They also naturally depend on the effector type, and may change for T cells.

The non-monotonic dependence of bridging molecules density with bsAb concentration is a new illustration of the known hook effect.^43, 45, 49^ While documented for *cis* binding, it is to our knowledge hardly considered when estimating effective concentration of cell engagers,^27^ which could considerably complicate the determination of the optimal quantity of bsAb to be administrated in a clinical setting. However, within the range of explored concentrations, the fits of individual lysis curves did not exhibit a significant hook effect, as shown on the residuals of Fig. S10A. Indeed, the linear approximation provides a satisfactory description for high HER2 density, because EC_50_ is much lower than the peak concentration 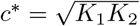(Fig. 2).

The multiscale model assumes that: (i) the molecular binding is at equilibrium at the synapse; (ii) the synaptic cleft has a uniform thickness; and (iii) the membrane receptors diffusion is negligible. While these assumptions are reasonable for SKBR3, which exhibits a high density of HER2 receptors, they may fail for MCF7-WT, which exhibits a low density of HER2. For example, an increase to the diffusion of HER2 towards the synapse of MCF7-WT could explain the lower values of *σ*_50_*/*Σ_2_ for certain donors compared to SKBR3. Also, the role of intercellular forces on binding/unbinding kinetics could become relevant at low bridging densities, where the cellular force is distributed over less bonds.^42, 47^ The topology of effector cell surfaces may also influence their lytic efficacy,^50^ which would require a refined description of the interplay between geometry and reaction. Another extension of our model could include multivalency effects in *cis* configuration on the effector or target side,^22^ in order to account for conventional bivalent Abs like herceptin, and the recent efforts to increase specificity of engagers *via* avidity.^51^

A practical consequence of the observed decoupling is that potency can be predicted, *for a given target cell line*, by the product of the medians by donors and bsAb, as shown on Fig. S9 and embodied in the parameters *α* and *σ*_50_ (Fig. S15). We also found very similar EC_50_ between two different cell lines exhibiting the same HER2 density: SKBR3 and MCF7-HER2+. The reduced potency *ec*_50_ explains quantitatively the dependence of EC_50_ on *K*_1_, *K*_2_, Σ_1_, Σ_2_. Therefore, affinities could be tuned independently, for example attenuated for clinical purposes,^14, 52^ while maintaining EC_50_ constant. Eq. 4 predicts the bsAb dose to selectively kill target cells based on their HER2 surface density. Interestingly, the model also sheds light on the behavior of conventional antibodies, like herceptin. For example, a low affinity of Fc fragment on the effector side can be compensated by an avidity effect caused by the two Fabs on the target side. In this manner, specificity (via the avidity) can be tuned while keeping a constant bridging potency. This was observed previously when comparing herceptin with bsAb.^38^

We show the usefulness of our approach to predict the potency of various engagers using Eq. 4 and compared to published data (Tab. 2). We also propose in Suppl. Mat. a practical protocole to predict the potency of new bispecific engagers.

The multiscale approach proposed here to describe a tripartite cell-molecule-cell system could also be relevant in various physiological or therapeutical situations where Fc receptors are involved, including when considering interactions of tumour cells with other immune cells, like macrophages, neutrophils, or den-dritic cells,^54^ as well as for T cell engagers. Our method and results can be exploited as general designing rules for therapeutic antibodies as well as for personalized medicine approches. Antibody engineering can be used to modulate the different affinities and number of valencies of immune cell engagers but can also tune other characteristics such as size, flexibility, and general geometry of therapeutic constructs, all of them expected to influence the cooperativity factor *α*. Future work will be needed to explore the possibility to rationally optimize this factor, opening the door to highly efficient therapeutics.

### Study limitations

Despite our confidence in our findings, we acknowledge limitations in our study that should be considered when trying to generalize the results. A first limitation comes from the reduced number of target cell lines, while the number of donors (15) and bispecific molecules (6) tested is rather extended compared to other published studies in the field. Comparing MCF7-WT and MCF7-HER2 has the purpose of assessing directly the effect of HER2 density Σ_1_, all other cell parameters being assumed to be constant. Additionally, comparing SKBR3 and MCF7-HER2 shows that different cell lines with the same level of HER2 expression behave very similarly. Addition of more target cell lines would have considerably increased the number of conditions tested (ca. 90 per cell line). Nevertheless, for use in a clinical setting, the best choice would be to use the patient target cells.

Our *in vitro* study on a 2D co-culture should help in accelerating antibody screening, but is not aimed at recapitulating the full complexity of bsAb-induced tumour cytotoxicity *in vivo*. Our theoretical modeling provides a quantitative prediction of bsAb in vitro potency based on a model, particularly on Eq. 4, which makes the contributions of the donor effector cells, the target cell line and the bsAb explicit. For such a model to be effective, some estimates of each parameter in the equation are required. While they can be determined by classical assays, minute differences can occur depending on the experimental setting. For example, Abs affinities can be determined on purified proteins or in cells, and using single nanobodies or bsAb. Table S2 shows that the differences are typically limited to a factor 3. Further differences in apparent affinities or avidities could be expected for multivalent engagers. Nevertheless, Eq. 4 should remain applicable in a pfirst approximation when using realistic avidities values.

Of note, the presented theoretical model was conceived for any engager, but its application was demonstrated only on NK cells. Nevertheless, our assumptions are consistent with the currently most used models for immune cell activation, like the kinetic segregation model, which is applicable for activating receptors from NK cells as well as T cells. We also propose consistent quantitative predictions from T cell engagers using data extracted from the literature. Finally, our study considers a fixed bsAb format, which helps to assess the role of other (molecular/cellular) parameters included in the model. An important conclusion is that the effect of the format should be contained in the *α* parameter, paving the way for future studies.

## Material and Methods

### Nanobodies and bispecific synthesis

Nanobodies anti-CD16 C21 and C28 were previously generated after immunization of llamas with the recombinant human Fc*γ*RIIIB and selected by phage display as described in.^41^ GenBank accession number are: EF5612911 for C21; EF561292 for C28.

Nanobodies anti-HER2 CA5, CE4 and C7b were previously generated after immunization of lamas with HER2-expressing SK-OV-3 ovarian cancer cells as described in.^55^ GeneBank accession numbers: JX047590 for C7b Production and purification of six bsAbs antiCD16xHER-2. These bispecific antibodies were generated by fusing single-domain antibodies targeting the human HER2 antigen (sdAb CE4, CA5 or C7b) and the activating FcgRIIIa receptor (sdAb C21, C28) to human Ck and CH1 IgG1 domains respectively. bsAb have 2 tag on CH1 (cmyc and 6HIS) and 1 tag on Ck (Flag).

Antibodies were transiently expressed in Expi293F cells (A14528, GIBCO) using the Expifectamine 293 Transfection kit (A14524, GIBCO) following the manufacturer’s procedure. Briefly, Expi293F cells were co-transfected with 30 µg of each pHLsec plasmids coding for the anti-HER2 Nanobody-Ck fusion and the anti-CD16 nanobody-CH1 fusion, in a 30 mL Expi293 expression medium at 37°C, 8% CO_2_. Proteins were harvested from the culture medium 5 days posttransfection. Cell-free supernatants were dialyzed against PBS overnight at 4°C and the His-tagged bsFabs/bsAbs constructs were purified by Immobilized Metal Affinity Chromatography on TALON superflow (Sigma) cobalt resin. The absence of aggregate was verified by size exclusion chromatography on a Superdex200 increase 10/300 column (Cytivia).

### Effector and target cells

Primary human NK cells from healthy donors were isolated from blood samples provided by the EFS (Marseille, France) by negative selection using the MACSxpress Whole Blood human NK cell isolation kit (Miltenyi Biotec cat.130-098-185), according to the manufacturer protocol. Purity of NK cells was determined by flow cytometry. Isolated primary NK were aliquoted into a 96-well round bottom plate (Corning) in RPMI 10% FBS at 2×10^5^ cells/well. Cells were centrifugated at 700 g for 2min at 4°C and medium was removed. Cells were incubated for 1h at 4°C and shaked at 700 g with 50 *µ*L of antibody: IgG1-PE (cat. 130-092-212) and antibody CD3-PE (cat. 130-091-374) as negative control. As positive controls, antibody CD56-APC (cat. 130-113-310) and antibody CD16-FITC (cat. 130-091-244) antibodies (all from Miltenyi Biotec) were used. Cells were stored in RPMI 1640 medium (Gibco, Life Technologies) complemented with 10 % foetal bovine serum (FBS, Gibco, Life Technologies) at 37 °C and used in the following 24h.

SKBR3 (ATCC HTB-30) is an epithelial cell line established from the mammary gland of a 43-year-old woman with adenocarcinoma in 1970. This cell line express naturally HER2 on cell surface. This cell line was used in previous studies to evaluate cytotoxic activity of bsFabs anti-HER2.^38^

MCF7-WT (ATCC HTB-22) is an epithelial cell line established from the mammary gland of an 69-year-old woman with adenocarcinoma. This cell line expresses naturally HER-2 on cell surface at low levels, since they do not have amplification of the HER2 (ErbB2) oncogene.^56^ This cell line was used in previous studies to evaluate cytotoxic activity of bsFabs antiHER2.^38^

MCF7-HER2+ is a genetically modified version of MCF7-WT which over-expresses HER2. To obtain them, MCF7 cells were electroporated with 1*µ*g of DNA plasmid HER2-GFP from Sino biologicals (ref HG10004-ACG) with Nucleofector 2b device (Lonza), and selected by antibiotic hygromycine. The expression of HER2 receptor was controlled by flow cytometry using LSRFortessa X20 (BD Biosciences, Franklin Lakes, NJ), using anti-human Her2 antibody (clone 9G6, ref sc-08, Santa Cruz Biotechnology, Dallas, Texas, RRID: AB627998). Cells expressing high level of HER2 receptor, were sorted and cloned with BD FACSMelody cell sorter (BD Biosciences, Franklin Lakes, NJ).

All target cells were cultured in RPMI 1640 with 10 % of FBS.

### Binding assay by flow cytometry

Primary NK cells were aliquoted into a 96-well round bottom plate (Corning) in RPMI 10% FBS at 2×10^5^ cells/well. Cells were centrifuged at 700 g for 2 min at 4°C and medium was removed. Titrations of the sdAbs or bsAbs were prepared in PBS/BSA 1% and added to primary NK cells. The range of tested concentration was typically between 0.1 nM to 1000 nM. As a negative control antibody human isotype control IgG1-PE (Biolegend 403504) was used. Cells were incubated with 50*µ*L of the primary antibody solutions for 1h at 4°C and shaked at 700 g followed by three washes in 200*µ*L of cold PBS/BSA 1%. The cells were resuspended in 50*µ*L of His-PE antibody (Miltenyi Biotec 130-120-718) and incubated for 1h at 4°C and shaked at 700 g. Following three washes in 200*µ*L cold PBS/BSA 1%, cells were analyzed on a MACSquant X flow cytometer (Miltenyi Biotec). K_*d*_ were determined by plotting the geometric median of signal versus log-concentration and using non-linear regression curve fitting using Prism software (GraphPad).

Her2 (SKBR3/MCF7) cells were aliquoted into a 96-well round bottom plate (Corning) in PBS+1%BSA at 2×10^5^ cells/well. Solution of 200nM of each bsAb is preincubated with 200nM anti-His Ab, for 30min at room temperature with shaking. Titrations of the premixed bsAb-anti His were prepared in PBS/BSA 1% and added to cells. As a positive control, a mouse monoclonal antibody anti-Her2 (clone 9G6) was used. As a negative control antibody human isotype control IgG1 (LEAF Purified Human IgG1 isotype control antibody, Biolegend 403502) was used. Cells were incubated with 100*µ*L of the antibody solutions for 1h at 4°C and shaked at 700 g followed by two washes in 200*µ*L of cold PBS/BSA 1%. The cells were resuspended in 100*µ*L of a secondary antibody, Goat anti-mouse-PE and incubated for 1h at 4°C and shaked at 700 g. Following two washes in 200*µ*L cold PBS/BSA 1%, cells were analyzed on a MACSquant X flow cytometer (Miltenyi Biotec). *K*_*d*_ were determined by plotting the geometric median of signal versus log-concentration and using non-linear regression curve fitting using Prism software (GraphPad).

### Cell receptors quantification

Quantification of CD16 on primary human NK cells was performed the day of the cell purification, by quantitative cytometry using a Biotin anti-human CD16 Antibody (clone 3G8; biolegend ref 302004), a Biotin Mouse IgG1*κ* as Isotype Ctrl Antibody (biolegend ref 400103) and the secondary antibody (mouse anti-IgG polyclonal FITC), as well as calibration beads from a commercial kit (CellQuant calibrator kit, ref 7208, Biocytex), used according to supplier’s recommendations.

Quantification of HER2 on tumour target cells (MCF7-WT, SKBR3 or MCF7-HER2+) was performed by quantitative cytometry using a anti-ErbB2/HER2 (clone 9G6) (Santa Cruz biotechnologies ref sc-08), a purified Mouse IgG1*κ* as Isotype Ctrl Antibody (Biolegend ref 400124) and the secondary antibody (mouse anti-IgG polyclonal FITC), as well as calibration beads from a commercial kit (CellQuant calibrator kit, ref 7208, Biocytex), used according to supplier’s recommendations.

### Luminescent assay for ADCC measurements

Target cells were plated at 5000 cells/well in 96-well plates (Greiner cat. 655083) and incubated at 37°C, 5% CO_2_ for 8h. Serial dilutions of bsAbs were added and cells were incubated for 30 min. Isolated primary NK cells were added (effector:target ratio of 5:1). Cells were further incubated overnight and the supernatent was rinsed. The cell viability was measured using CellTiter-Glo reagent (Promega), according to the manufacturer’s instructions. The assay reports the amount of ATP from living cells in the sample, which includes living targets as well as living attached NK. An analysis was developed to determine the contribution to the ATP luminescence signal of living effector cells bound to their targets. For this, NK cells were preloaded with 1*µ*g/ml CFSE. A calibration of fluorescence signal as function of the number of primary NK effector cells was performed by measuring variable amounts of NK (Fig. S18A). This relation was fitted as follows:

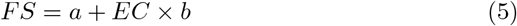

where FS is the fluorescence signal of the sample, a the *y*-intercept and b the slope of the linear fit of the fluorescence calibration. The ATP signal that corresponds to the effector cells (ES_*AT P*_) was determined using the following luminescence calibration

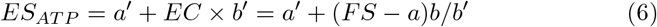

with a’ and b’ previously calibrated, as shown (Fig. S18B). By removing this ES_*AT P*_ from the total ATP signal (TS), the target cell ATP signal (TS_*AT P*_) was determined.

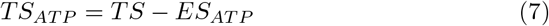

Once TS_*AT P*_ is known and using the ATP signal of target cells alone as live target cells control (T_*live*_), the fraction of target cells lysis was determined.

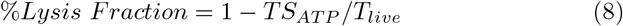

### Real-time cytotoxicity assay

The cytolytic potential of isolated primary NK was analyzed in a real-time cytotoxicity assay with an xCELLigence RTCA SP instrument (ACEA Biosciences, San Diego, CA, USA).^57^ In each well 4*×* 10^4^ MCF7-HER2+ cells were seeded. When Cell index (CI) were close to 1, dilutions of bsAbs (100pM, 10pM and 1pM) and 2 *×*10^5^ primary NK cells were added for an effector:target ratio of 5:1, neglecting the proliferation of target cells. Cell viability was monitored every 10 min for typically 24h. Cell indexes (CIs) were normalized to CI of the time-point before bsAbs and primary NK cells addition and specific lysis was calculated in relation to the control cells lacking any effector primary NK cells, following Eq. S6.

## Author Contributions

LL, PC, FP and PR designed the research. CG, AA, MB carried out experiments. AA, PC, BK and CG contributed reagents. LL and FP developed the model. CG, PHP, PC, FP and LL analyzed data. LL, FP, PC wrote the manuscript. All authors critically revised the manuscript.

## Acknowledgements

We thank A. Raguin for careful reading of the manuscript. This work was supported by AMIDEX Emergence Innovation (project ForSelecAntibody) and Plan Cancer PhysCancer program (project ComPhysAb).

## Supporting Information Available

Additional methods, analyses and figures in supplementary information

## Supplementary Material

### Analysis of bsAb and donor ranking by the potency

The ranking based on EC_50_ is quantified by comparing the donor (respectively the Ab) order, as gauged by EC_50_ for each possible pair of Ab (respectively the donor) (data from Fig. 1). More precisely, when ranking by bispecific Ab, we used a pair of donors and evaluate the Spearman coefficient between the two series of EC_50_ values found for all the considered bsAbs. If the ranking of EC_50_ is exactly the same for both donors, this particular pair will display a Spearman coefficient *by bsAb* of 1. We repeat this procedure for all possible pairs of donors and then build the cumulative distribution of the ensuing Spearman coefficients, i.e. the fraction of coefficients found to be lower or equal than a given value (between 1 and 1). We follow a similar procedure when ranking EC_50_ values *by donor* for given pairs of bsAb.

We illustrate this comparison in Fig. S4. For example, one can read immediately from panel A that only about 5 % of all donor pairs feature EC_50_ series (corresponding to different bsAbs) that share less than 60 % of their bsAb rankings. Similar plots for Min and Max parameters are reported in Fig. S4B,C, showing no dependence of Min on bsAb, and a mimimal dependence of Max on bsAb. This corresponds to curves close to the diagonal, (dashed line in Fig. S4). On the other hand, both Min and Max exhibit a strong dependence on the donor. The areas under the cumulative curves can be considered as a measure of the deviation from randomness. In fact, for infinite-length random vectors, the probability density of pair Spearman coefficient is a delta function at zero correlation, and thus the cumulated fraction is a Heaviside theta function, equal to 0 for negative coefficients and equal to 1 for positive coefficients. Consequently, the areas under the curves in the (P(correlation*<*S),S) plane are equal to 0 for random orderings and equal to 1 for perfect ranking correlation among all vectors. The areas measured for our samples are reported in Table S3, where the values close to 1 that indicate a strongly conserved order are highlighted in bold.

### Model for bispecific antibody binding at equilibrium

The aim is to predict the surface density *σ* of bridging bsAb at the target-effector synapse at equilibrium. The reaction scheme is shown in Fig. 2A and notations are explained in Table 1. 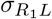 and 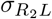 are the surface densities of *R*_1_*L* and *R*_2_*L* respectively. Combination of Eqs. S1a and S1c from Tab. S4 leads to Eq. S1.

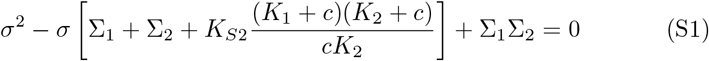

We have checked that in our experimental conditions there is no significant depletion of antibody in the sample volume upon reaction on the cell surfaces.

By applying the detailed balance condition to the reaction cycle (zero net probability current along the cycle at equilibrium), one obtains a simple relation that constrains the different dissociation constants, namely

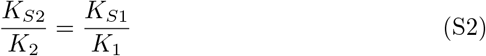

which has been already noticed in the case of multivalent molecules binding to membrane receptors on the *same* membrane.^58^ In view of this kinetic scheme, it is natural to introduce a dimensionless cooperativity parameter, defined as

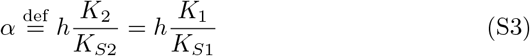

where *h* is the intermembrane distance at the synaptic cleft. The cooperativity *α* represents the factor of gain (*α*≫ 1), or loss (*α*≪ 1) of affinity upon binding in the second step to one of the surfaces when one bond is already present at the opposite surface, as compared to binding in the first step to the same surface directly from the solution. A similar cooperativity parameter was previously proposed to describe divalent ligand-monovalent molecule binding.^44^ With this notation, Eq. (S1) can be symmetrized as

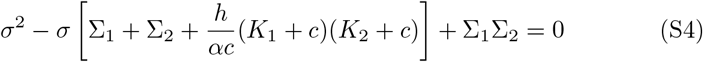

### Cell population dynamics

*At the cell population scale*, the variation with time of the number of living target cells, *T* (*t*), depends on the number of effector cells *N*_0_, the total lysis rate *k*_*a*_ and on the fraction of active effector cells, which we term *γ*(*t*). Note that target cells may still proliferate with a rate *k*_*p*_.

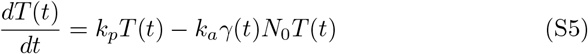

The above equation can be readily integrated as

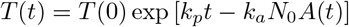

where 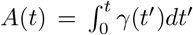 with *A*(0) = 0. The total lysis at time *t* is defined as the complement to the survival probability corrected for the proliferation of target cells, namely

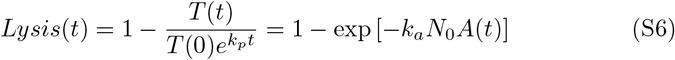

### Model-free decoupling of bsAb and donor contribution to potency

In the general case, the decoupling of the contributions of bsAb *i*, donor *j*, and target cell line *k* to EC_50_ can be expressed by writing it as a product of 3 factors: 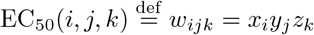.An estimator of EC_50_(*i, j, k*) is then:

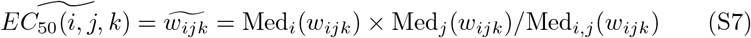

where Med_*n*_() takes the median value of *w*_*ijk*_ on index n. As shown on Fig. S9, *w*_*ijk*_ and 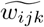 exhibit a good correlation for each target cell line *k*=1,2,3, with Pearson and Spearman coefficients higher than 0.9. This observation is supporting the two hypotheses in the main text, which postulate a decoupling between the contributions of bsAb and donor to EC_50_, via *α* for bsAb and *σ*_50_ for donors. Formally, considering the target cell line *k*, the bsAb i dependent term is Med_*j*_(*w*_*ijk*_) (median taken on all donors) and the donor j dependent term is Med_*i*_(*w*_*ijk*_) (median taken on all bsAbs).

### Exploitation of the reduced potency *ec*_50_

*ec*_50_ scales as surface density containing the information on the donor through *σ*_50_ and on the bsAb through *α*. Those dependences can be quantified, without the need to rely on the results of the global fit of lysis curves by the multiscal model, by calculating the Spearman ranking by bsAb or by donor (see Fig. S19 and Tab. S3). The pairwise Wilcoxon coefficients were also calculated to compare bsAb and donors (Fig. S20). Significant differences on *ec*_50_ are observed on MCF7-WT for CA5-21 and CA5-28 with the other bsAbs (Fig. S20A). On both target cell lines, the differences mostly vanish for *ec*_50_*α* between bsAbs (Fig. S20C-D) or for *ec*_50_*/σ*_50_ between donors (Fig. S20G-H). The relative values of the fitting parameters *σ*_50_ and 1*/α* correlate well with the relative median of *ec*_50_, respectively calculated by bsAb and by donor (Fig. S15).

The correlation observed in Fig. 3C suggests an alternative way to estimate the parameters *σ*_50_ and *α*, namely by fitting them simultaneously so that *c*_50_ matches the experimental values of EC_50_ (from Hill analyses) for all the conditions considered. By doing this, we obtain a fair correlation between the results of a global fit of the lysis curves and those obtained by adjusting *ec*_50_ to EC_50_ in the case of SKBR3 (Fig. S21). However, there remains an underestimation of *α* obtained directly from EC_50_, that we attribute to the linearization. We also get a good match between *ec*_50_ and the quotient *σ*_50_*/α* obtained from EC_50_, as shown in Fig. S22.

### Estimate of the potency of immune cell engagers

We propose in this section a detailed protocol to estimate the potency of immune cell engagers. This method is applicable within the limitations stated at the end of the Discussion section. We are therefore considering a bispecific molecule (2 antibody specificities) targeting a tumour associated antigen (side 1) and an activating receptor of a immune effector cell (side 2, typically CD16 for NK or TCR-CD3 for T-cells). We use the approximation provided by Eq. 4 and corresponding parameters should be estimated.

1. Chose a representative target cell line and primary effector cell type.
2. Measure apparent affinities *K*_1_ and *K*_2_ on target and effector side respectively. Ideally, the measurement is performed by flow cytometry on each cell type using the engager and a secondary antibody (eg. anti-his if the engager possess a his-tag); refer to Method section “Binding assay by flow cytometry” and results in Table S2. Alternatively, data measured on recombinant proteins for each antibody specificity could be used.
3. Measure the surface density of receptors Σ_1_ and Σ_2_ on each cell type; refer to Method section “Cell receptor quantification”. Alternatively, refer to Tab. S1 or literature.
4. the donor sensitivity *σ*_50_ can vary between 0.1 molec/*µ*m^2^ (high responsive donor) and 10 molec/*µ*m^2^ (low responsive donor) (see Fig. S11B). In absence of any information about the donor, a median value can be taken within this range, typically 1 molec/*µ*m^2^. Note that this value determined here for NK cells may vary for T cells.
5. the bsAb cooperativity *α* = 0.02 is estimated for the bsFab format used in this study (Fig. S11A). While a different format may yield a different value for *α*, we show in the discussion that this value applies also for the popular format of knob and hole for T-cell engagers. The immune synapse thickness is fixed to *h* = 10 nm (Tab. 1).
6. If all parameters *K*_1_, *K*_2_, Σ_1_, Σ_2_, *α, σ*_50_ have estimates, Eq. 4 can be used to predict the potency. Alternatively, if some parameters are unknown, for example *α* corresponding to a different Ab format, some ratio of potency can be estimated based on the ratio of known parameters, as shown in Tab. 2.

## Supplementary tables

**Table S1:** Cellular and Receptor Data. Measurements of number of receptors were performed by flow cytometry using the CellQuant calibrator kit (see Methods). Values indicate mean *±* standard deviation.

| Receptor | Cell type | Diameter<br>($\mu\text{m}$) | Nb. Rec.<br>($\times 10^3$) | Density<br>(molec/$\mu\text{m}^2$) | Symbol |
| --- | --- | --- | --- | --- | --- |
| HER2 | MCF7-WT | $20 \pm 4$ <sup>59</sup> | $9 \pm 3.5$ | $7 \pm 3$ | |
| | SKBR3 | $18 \pm 4$ <sup>60</sup> | $615 \pm 100$ | $600 \pm 100$ | $\Sigma_1$ |
| | MCF7-HER2+ | $20 \pm 4$ | $617 \pm 178$ | $490 \pm 140$ | |
| CD16 | Primary NK | $6.5$ <sup>61</sup> | $40 \pm 10$ | $300 \pm 75$ | $\Sigma_2$ |

**Table S2:** Bispecific Antibodies Affinity Data. Apparent affinities were measured by flow cytometry (see methods). Values indicate mean*±* standard deviation.

| Receptor | Cell type | Bispecific | Affinity (nM) | Symbol |
| --- | --- | --- | --- | --- |
| HER2 | SKBR3 | CA5-21 | $10.7 \pm 1$ | $K_1$ |
| | | CE4-21 | $8.9 \pm 1.3$ | |
| | | C7b-21 | $27 \pm 1.9$ | |
|  | MCF7-WT | CA5-21 | 3.4 |  |
|  |  | CE4-21 | 3.6 |  |
|  |  | C7b-21 | 70 |  |
|  | MCF7-HER2+ | see SKBR3 |  |  |
| | Primary NK | CA5-21 | $11 \pm 2$ | $K_2$ |
| | | CA5-28 | $50 \pm 6$ | |

**Table S3:** Ranking of bsAb and donor using parameters from Hill or the multiscale cytotoxicity model, quantified by the areas under the curves representing Spearman vs pairs cumulative fraction in Figs. 1D, 1E, S4 and S19. The pool of 3 target cell lines are considered in each case. Values ≪ 0.5 indicate a random order, while values ≫ 0.5 (underlined) indicate a strongly conserved order.

| Model | Parameter ranked | Ranking by bsAb | Ranking by donor |
| --- | --- | --- | --- |
| Hill | $EC_{50}$ | <u>0.85</u> | <u>0.63</u> |
| Hill | Min | 0.01 | <u>0.83</u> |
| Hill | Max | 0.30 | <u>0.75</u> |
| Multiscale | $ec_{50}$ | 0.45 | <u>0.76</u> |
| Multiscale | $ec_{50}\alpha$ | 0.16 | 0.85 |
| Multiscale | $ec_{50}/\sigma_{50}$ | 0.48 | -0.03 |

**Table S4:** Equilibrium reactions and equations for bispecific binding at the immune synapse interface. See reaction scheme in Fig. 2A and notations in Tab. 1.

| Reaction | Dissoc. Const. | Detailed balance condition |  |
| --- | --- | --- | --- |
| $R_1L \rightleftharpoons R_1 + L$ | $K_1$ | $\sigma_{R_1L} = \frac{c}{K_1+c}(\Sigma_1 - \sigma)$ | (S1a) |
| $R_2L \rightleftharpoons R_2 + L$ | $K_2$ | $\sigma_{R_2L} = \frac{c}{K_2+c}(\Sigma_2 - \sigma)$ | (S1b) |
| $R_1LR_2 \rightleftharpoons R_1L + R_2$ | $K_{S2}$ | $\sigma K_{S2} = \sigma_{R_1L} \frac{K_2}{K_2+c}(\Sigma_2 - \sigma)$ | (S1c) |
| $R_1LR_2 \rightleftharpoons R_2L + R_1$ | $K_{S1}$ | $\sigma K_{S1} = \sigma_{R_2L} \frac{K_1}{K_1+c}(\Sigma_1 - \sigma)$ | (S1d) |

## Supplementary figures

**Figure S1:**
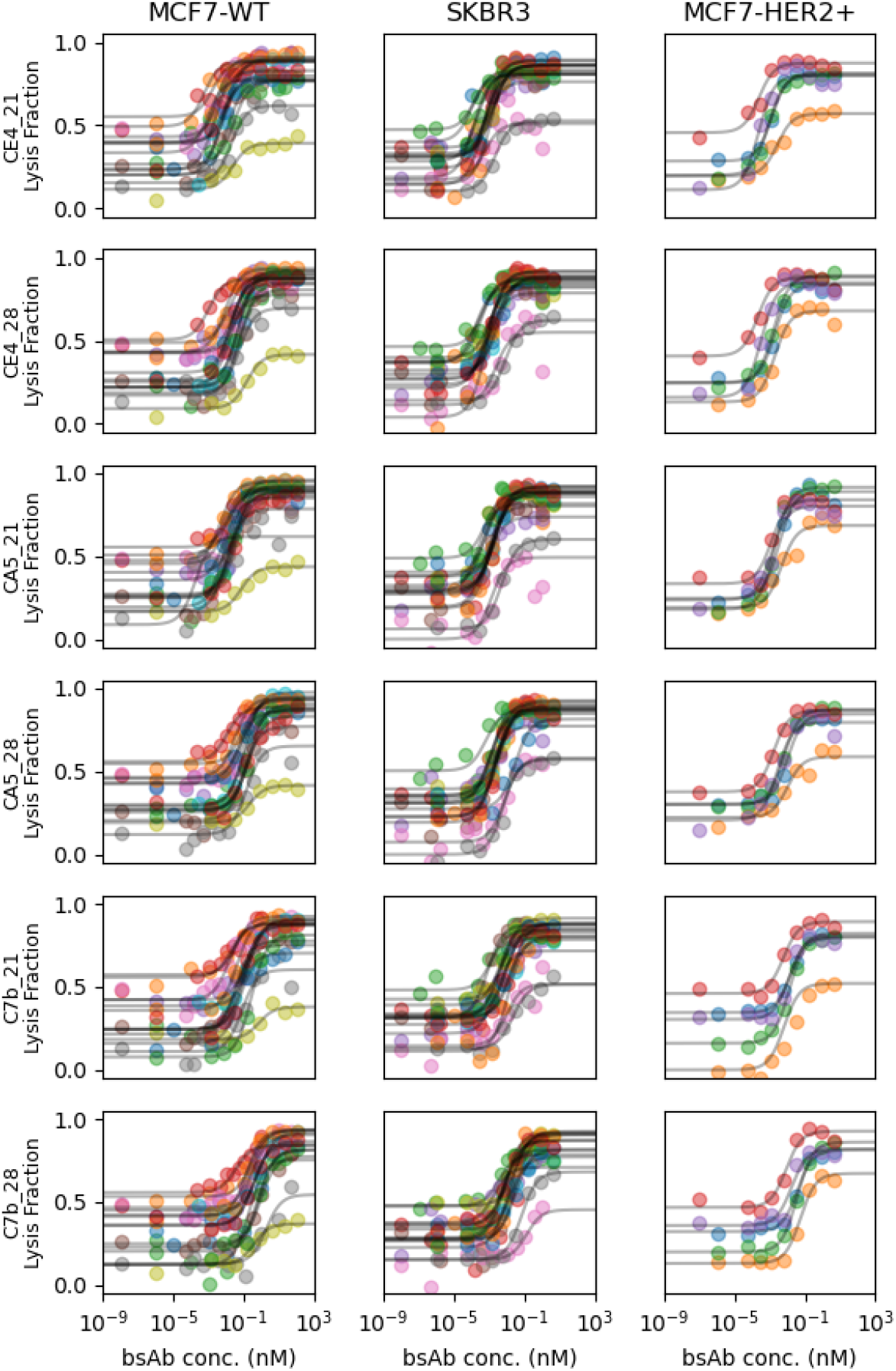
Ensemble of lysis measurements as a function of bispecific concentration obtained with luminescent cell viability assay. The series of curves are sorted by target cell line (column) and by bispecific (row). Within one graph, each color represent data of one NK cell donor. The solid lines are a Hill fit of the data using Eq. 1.

**Figure S2:**
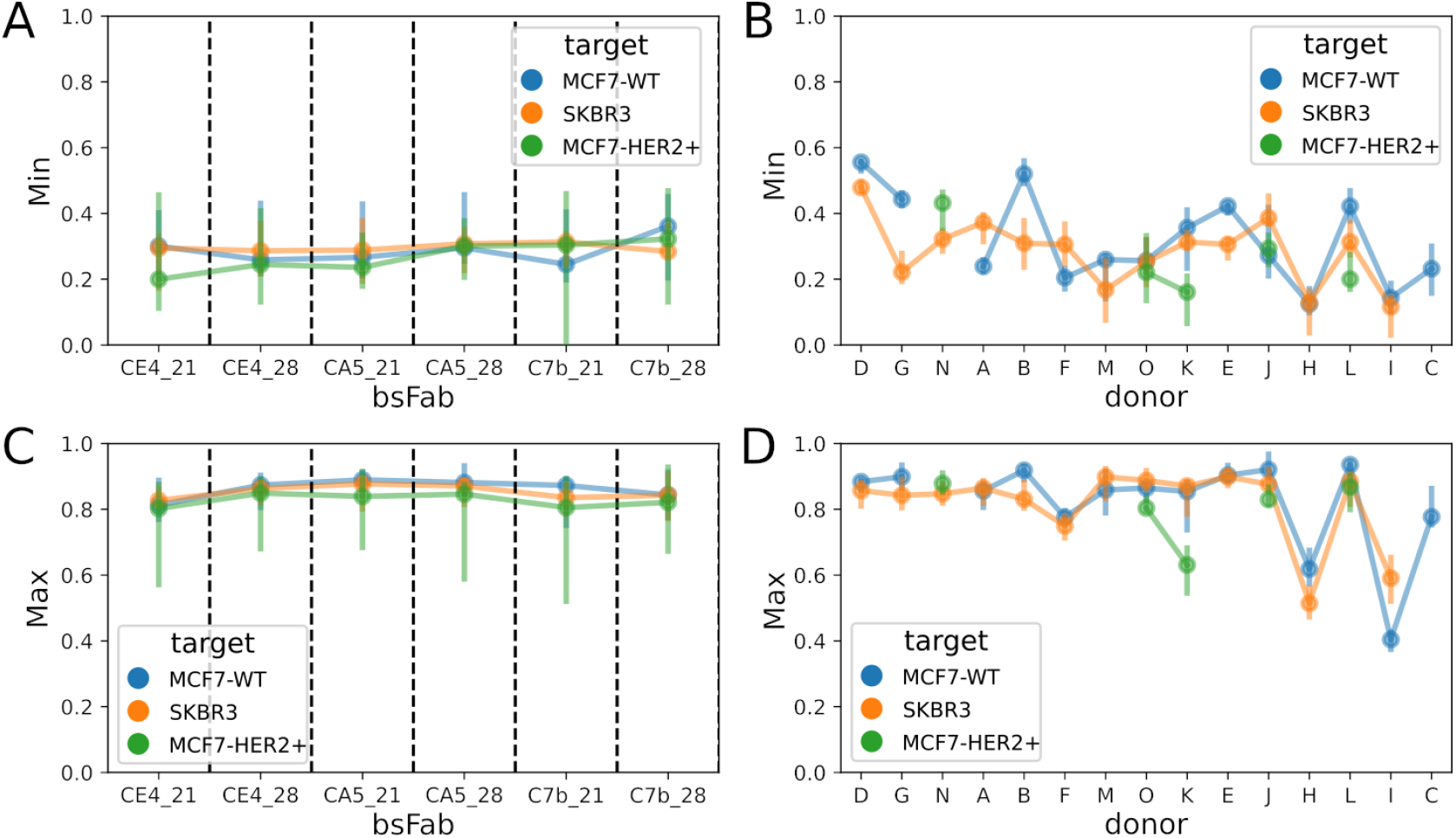
Hill fit parameters Min (A, B) or Max (C, D) sorted by bsAb (A, C) or by donor (B, D). Each point is the median on the donor (A, C) or the bsAb (B, D), with the bar representing 95% percentile interval. bsAb or donors are ranked according to the median value of EC_50_ for SKBR3 cell line.

**Figure S3:**
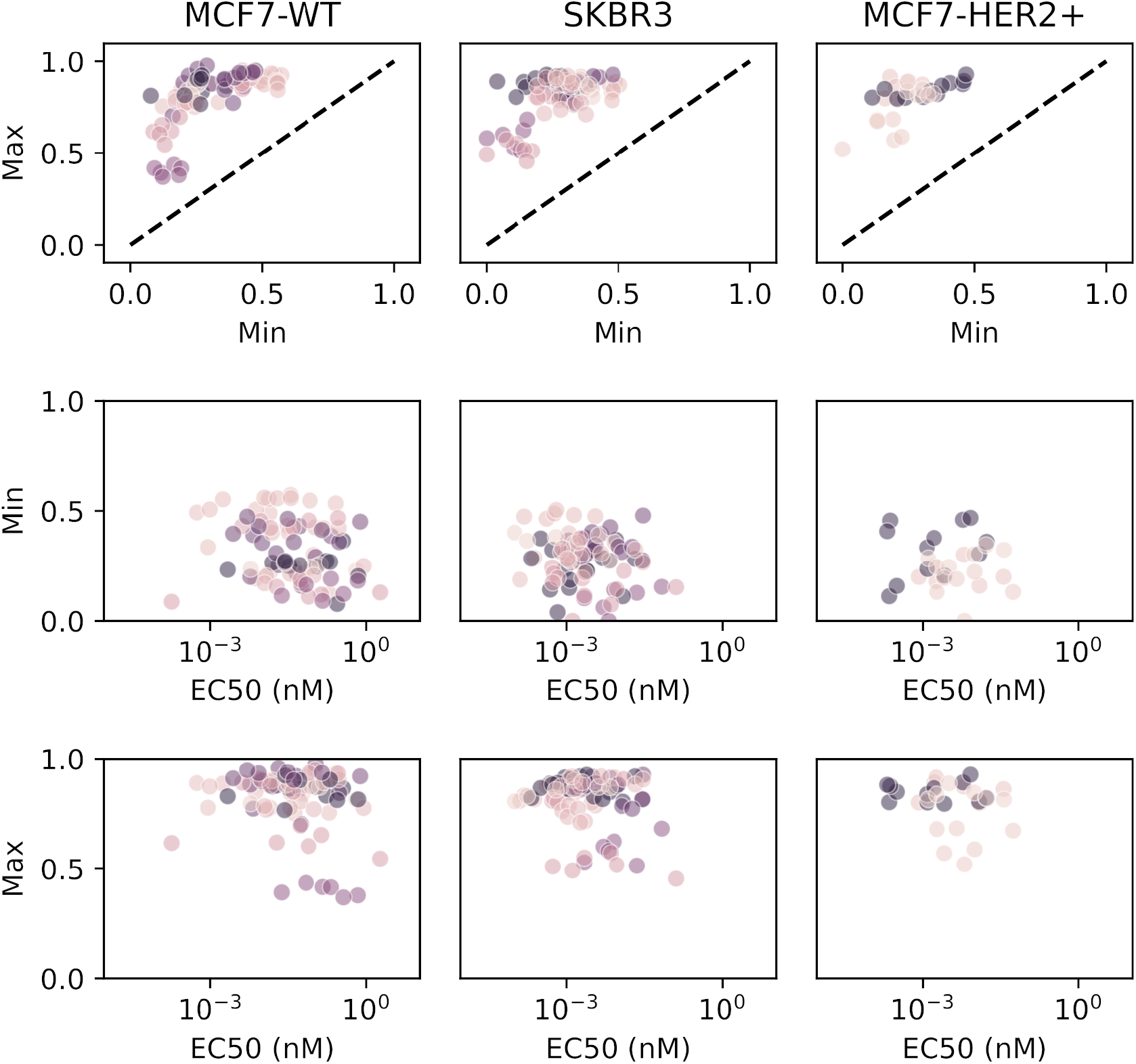
Relation between Hill parameters measured on individual *Lysis* vs *c* curves, shown separately for each target cell line. Each dot represents one biological condition (donor and bsAb). Dashed lines indicate *x* = *y*.

**Figure S4:**
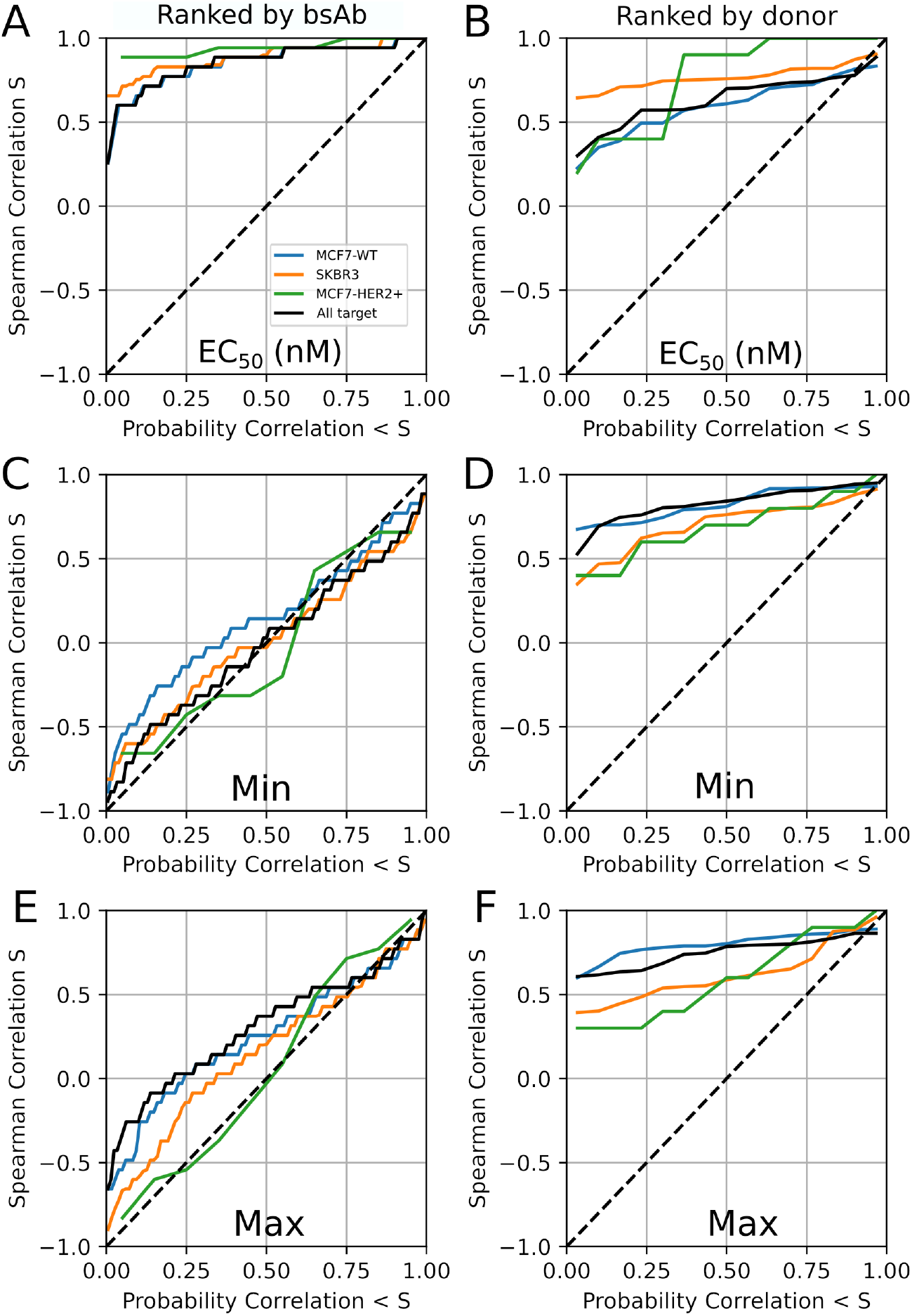
Spearman ranking based on Hill parameters for lysis EC_50_ (A,B) minimum lysis Min (C,D) or maximum lysis Max (E,F). A, C, E: ranking of bsAb for pairs of donors. B, D, F: ranking of donors for pairs of bsAb.

**Figure S5:**
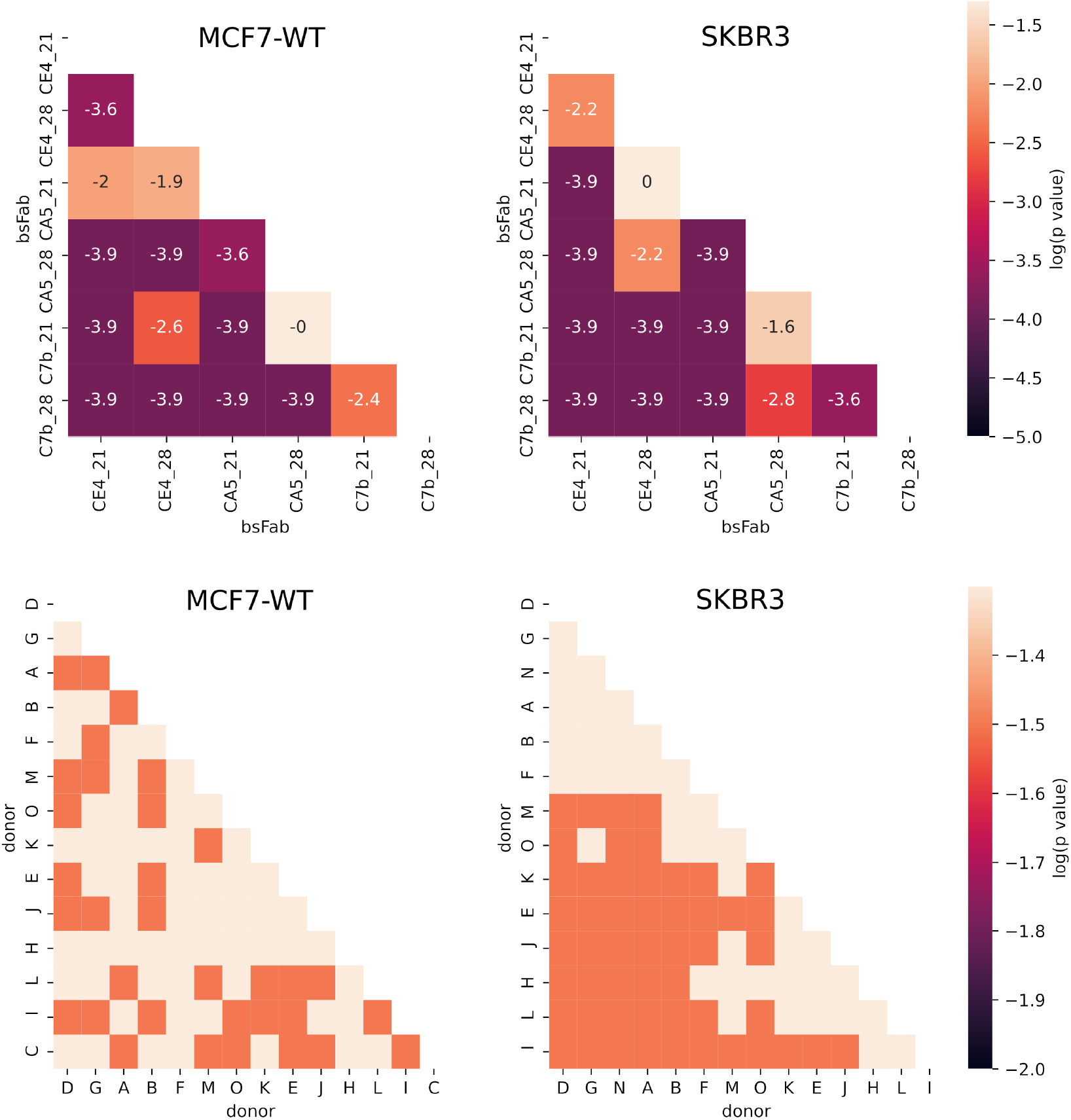
Matrix of p-values of paired Wilcoxon tests for differences in EC_50_ between bsAb (top row) or between donors (bottom row). bsAbs and donors are ranked by the median values corresponding to SKBR3 target.

**Figure S6:**
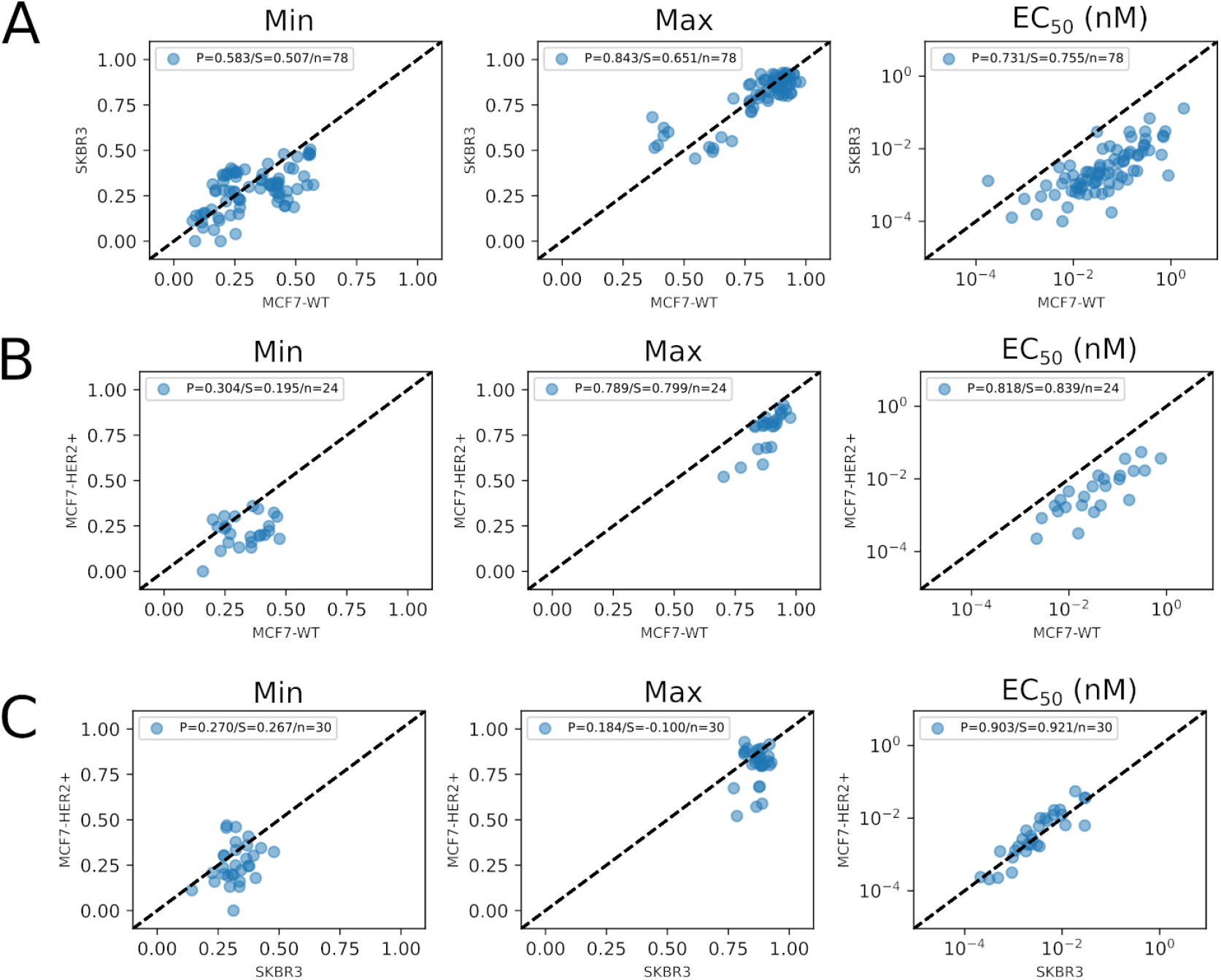
Correlation of Hill parameters per donor between target cell lines: MCF7-WT and SKBR3. B. MCF7-WT and MCF7-HER2+. C. SKBR3 and MCF7-HER2+. The dashed lines represent *x* = *y*. P: Pearson coefficient. S: Spearman coefficient. n: number of points.

**Figure S7:**
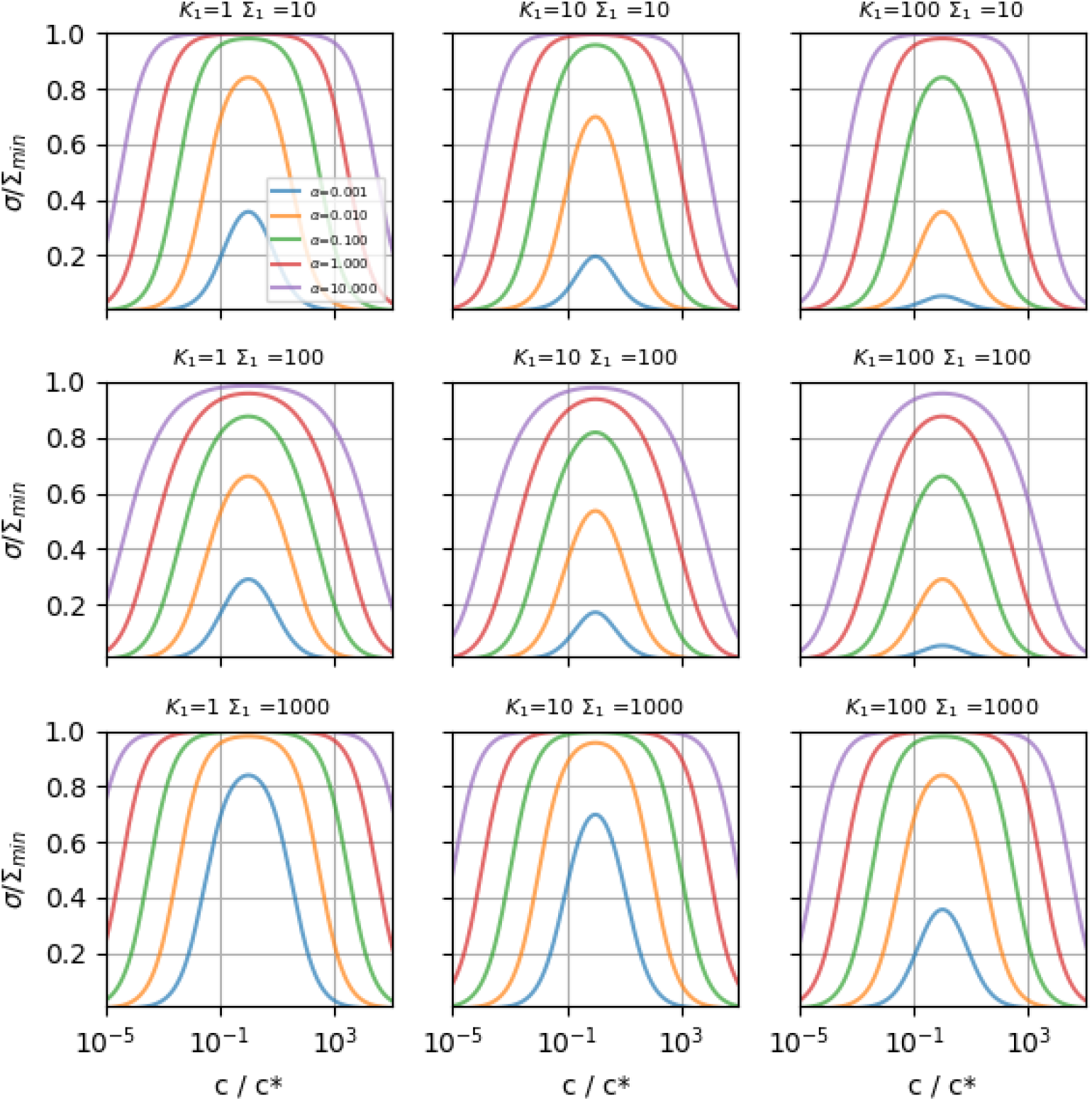
Exploration of density *σ* of bridging bsAb. Different columns are different values of *K*_1_ (in nM), while different rows correspond to different values of Σ_1_ (in molec/*µ*m^2^). *K*_2_ = 10 nM and Σ_2_ = 300 molec/*µ*m^2^ are fixed. *α* is varied and indicated as a color. Σ_min_ =min(Σ_1_, Σ_2_). 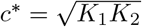.

**Figure S8:**
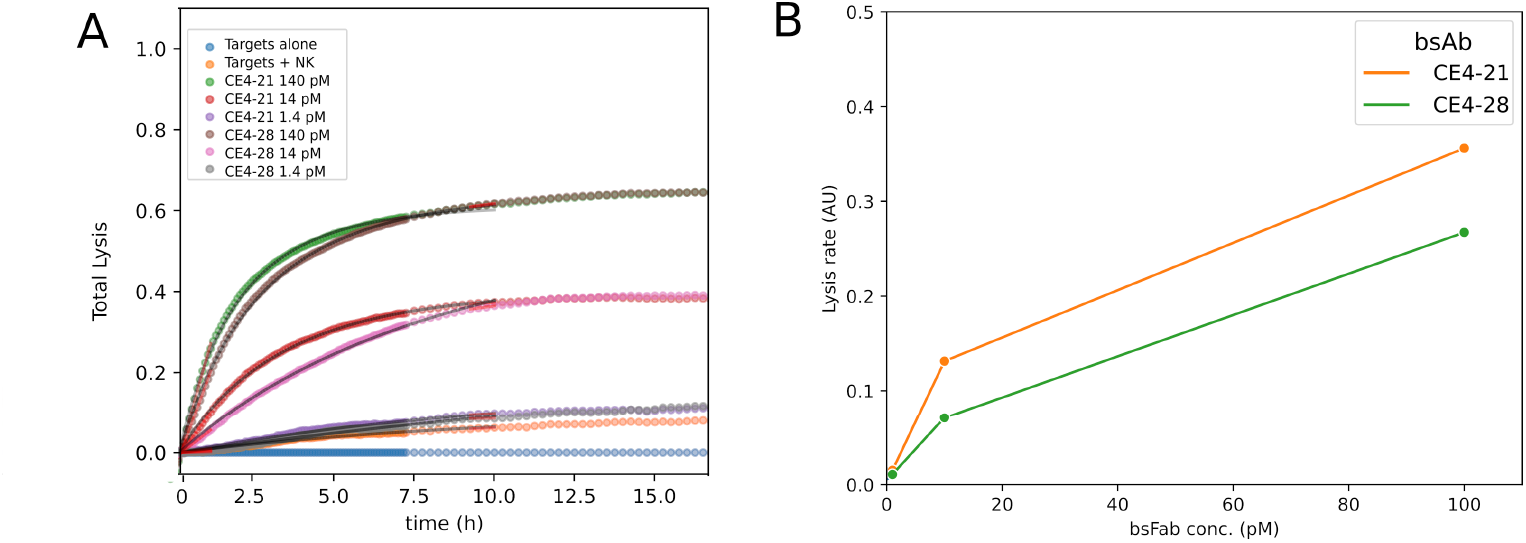
Typical real time cytotoxicity data. A. RTCA curves after normalization by the proliferation and substraction of the spontaneous lysis. The curves are fitted with Eq. S6, taking for *γ*(*t*) an exponential decay on 0.2 / hour. B. Measured initial lysis rate as a function of bsAb concentration.

**Figure S9:**
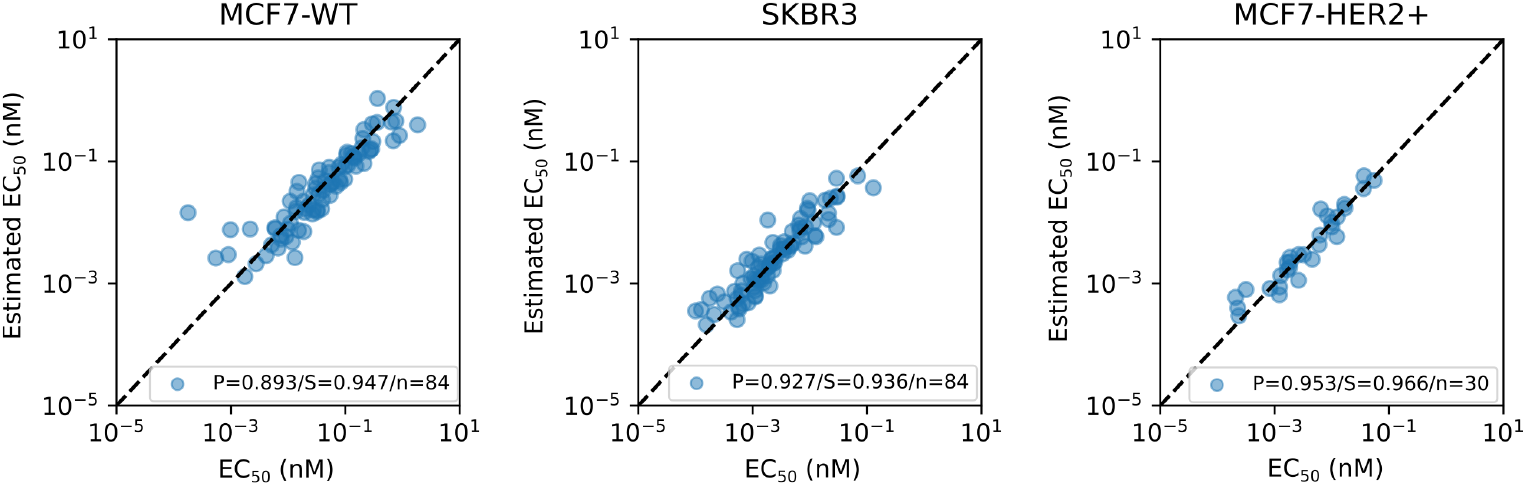
EC_50_ can be estimated from decoupled bsAb and donor contributions. Estimated EC_50_ is calculated from Eq. S7, separately for each target cell line, and plotted as a function of measured EC_50_. Each point represent one Donor/bsAb/Target condition. The dashed lines represent *x* = *y*. P: Pearson coefficient. S: Spearman coefficient. n: number of points.

**Figure S10:**
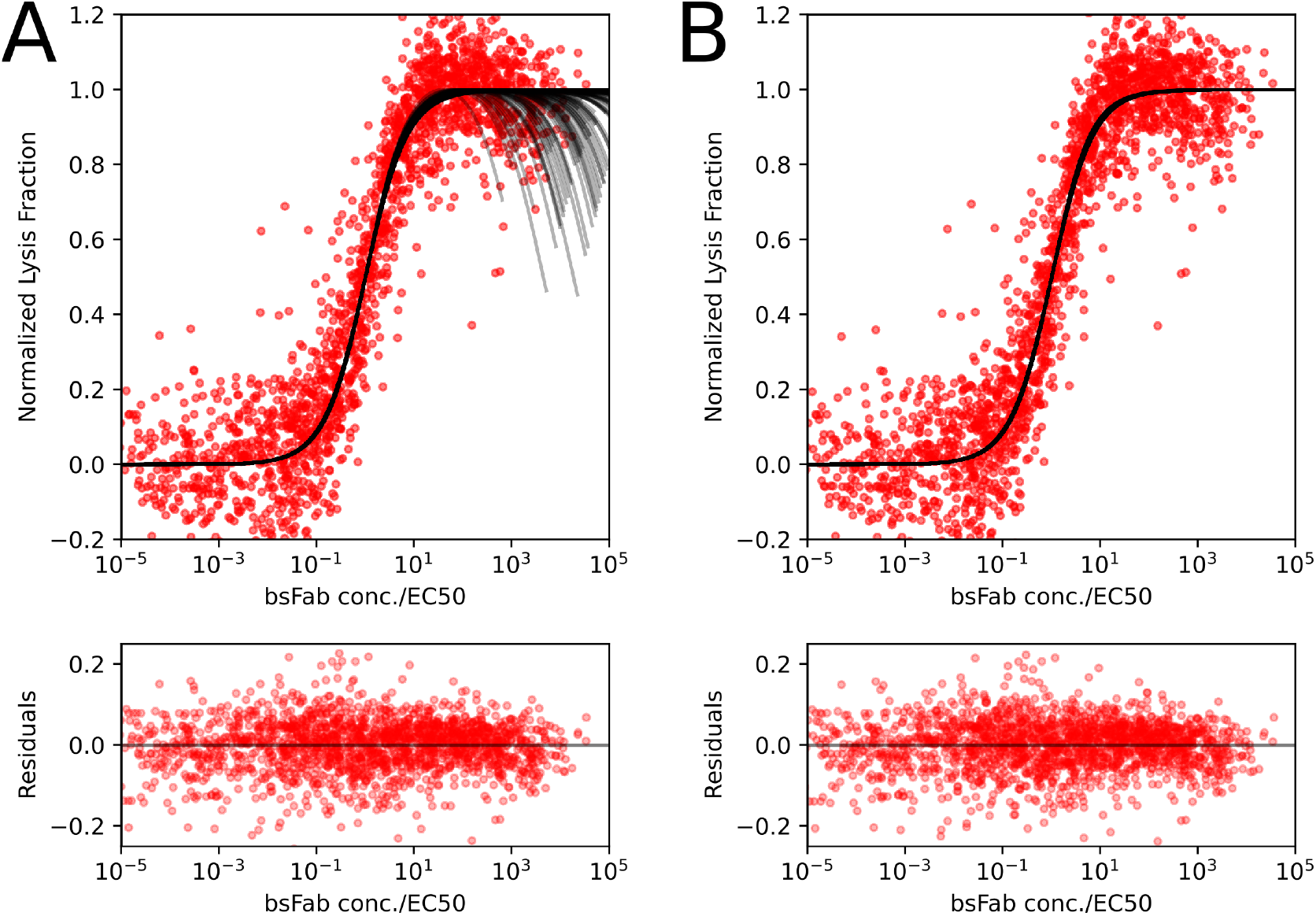
Normalized lysis curves using parameters from the multiscale model. Red dots: data. Black curves: fit. A. Non-linear version of the model with surface density of bridging solution of Eq. S4. B. linear version of the model with surface density of bridges 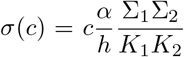.Each red dot represents one Donor/bsAb/Target condition. 198 conditions are represented.

**Figure S11:**
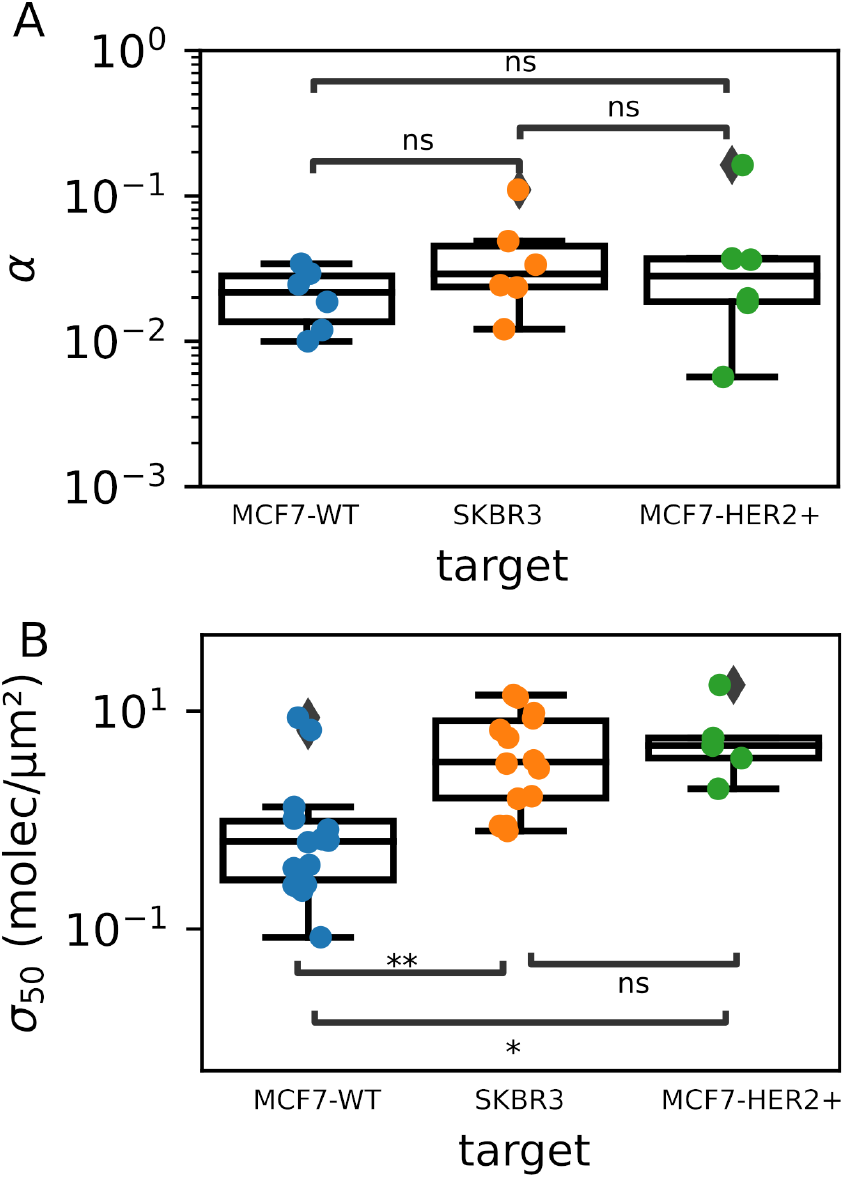
bsAb cooperativity (A) and NK threshold response (B) fitted from lysis data averaged over target cell line. The test is Mann-Whitney from Scipy library. *: P*<*0.05. **: P*<*0.01.

**Figure S12:**
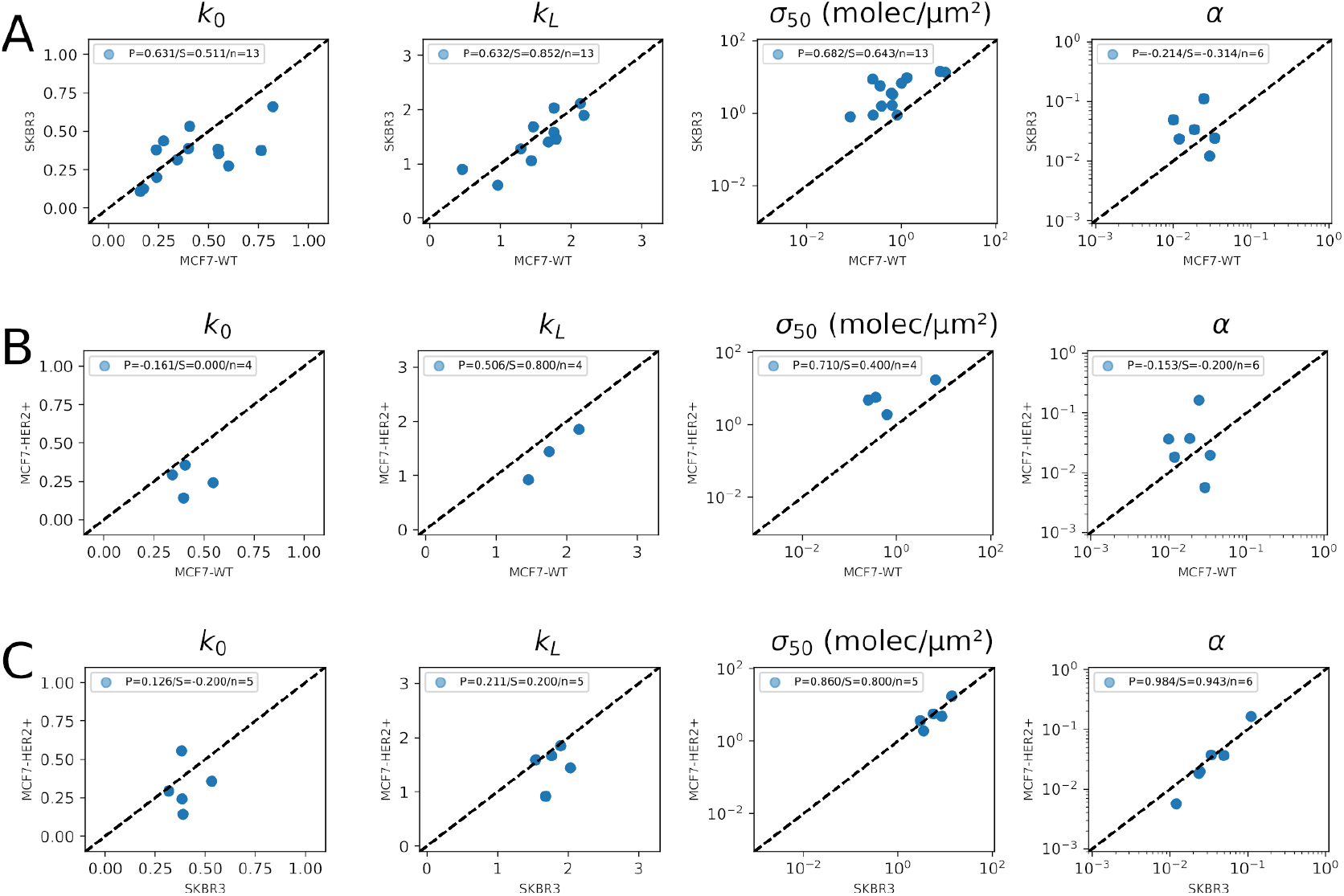
Correlation of best-fit parameters of the multiscale model between cell lines. A. MCF7-WT and SKBR3. B. MCF7-WT and MCF7-HER2+. C.SKBR3 and MCF7-HER2+. P: Pearson coefficient. S: Spearman coefficient. n: number of points. Dashed lines indicate *x* = *y*.

**Figure S13:**
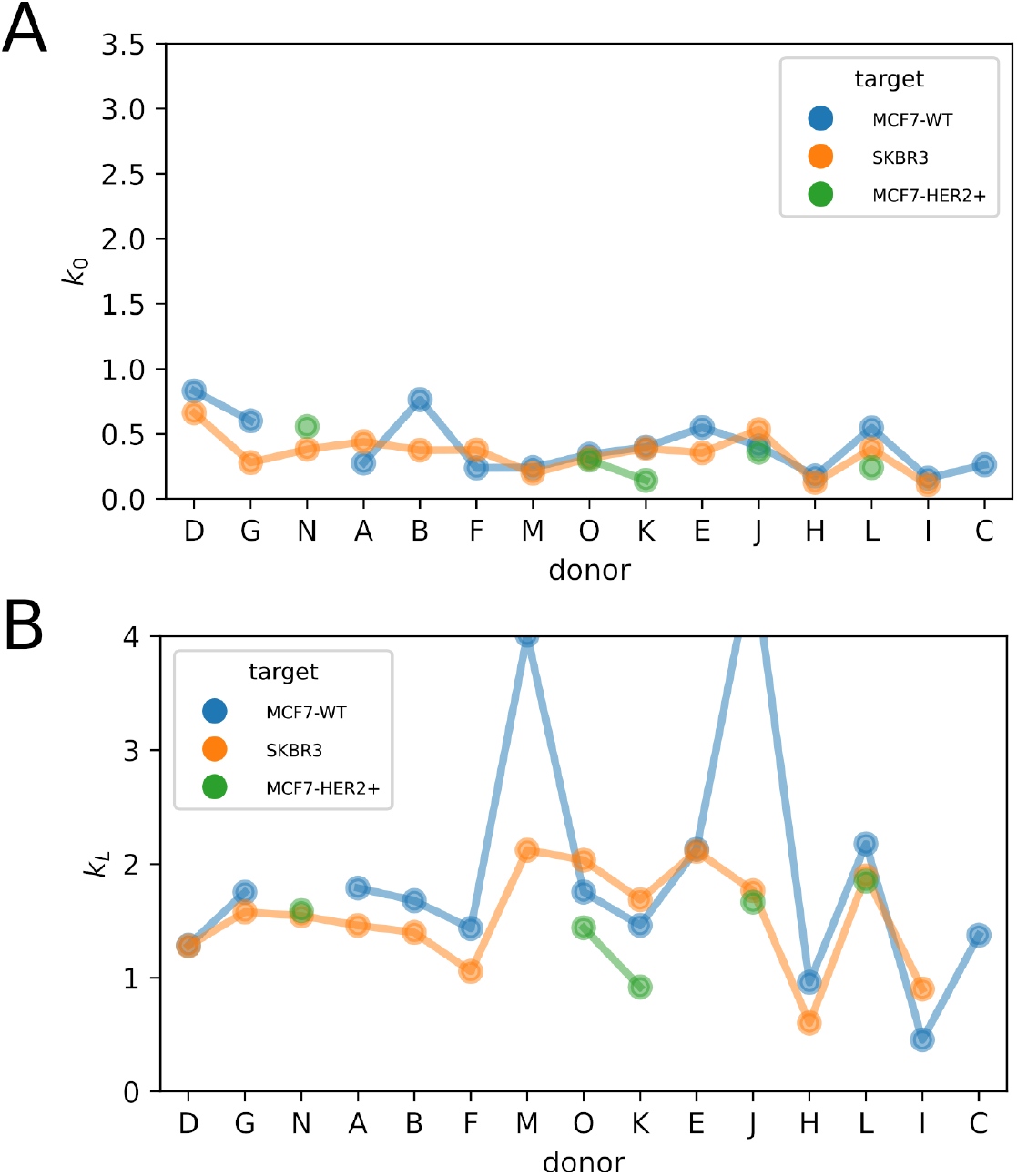
Lysis rate parameters for spontaneous lysis *k*_0_ (A) and ADCC *k*_*L*_ (B) obtained from fitting the complete dataset. Units are *N*_0_*A*(*T*), see Eq. S6.

**Figure S14:**
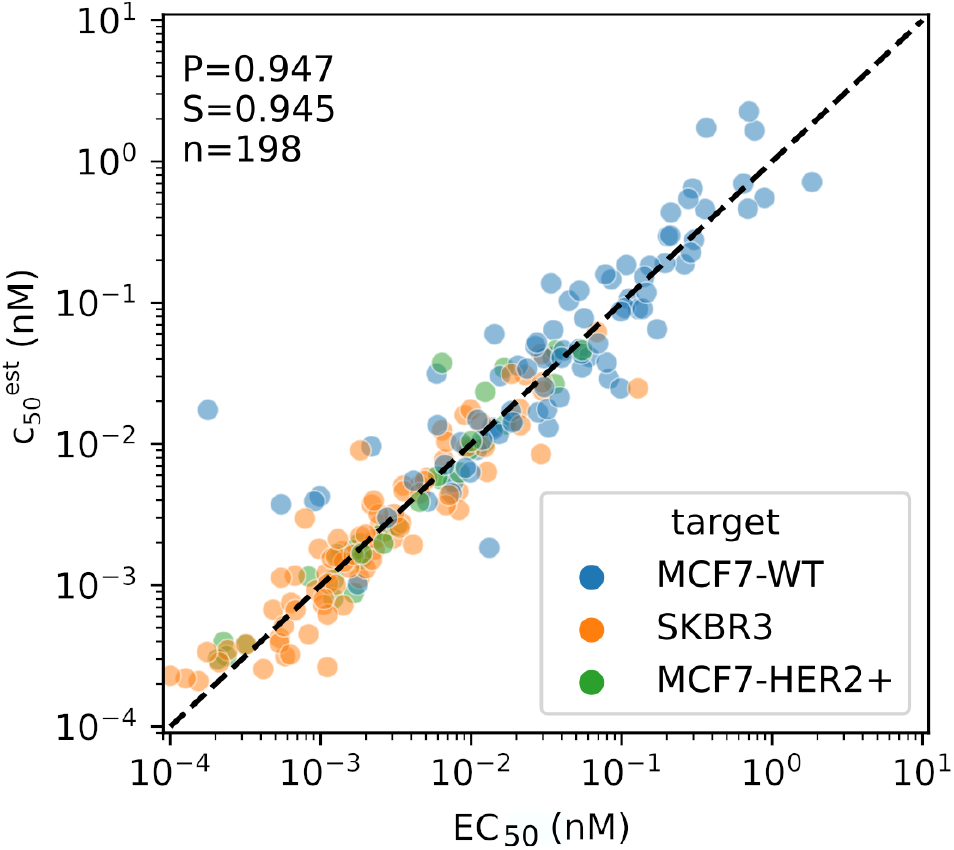
Refined estimate of EC with 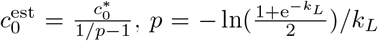 and 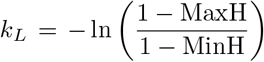. Each point represents one Donor/bsAb/Target 1 – MinH condition. P: Pearson coefficient. S: Spearman coefficient. n: Number of points . Dashed line indicates *x* = *y*.

**Figure S15:**
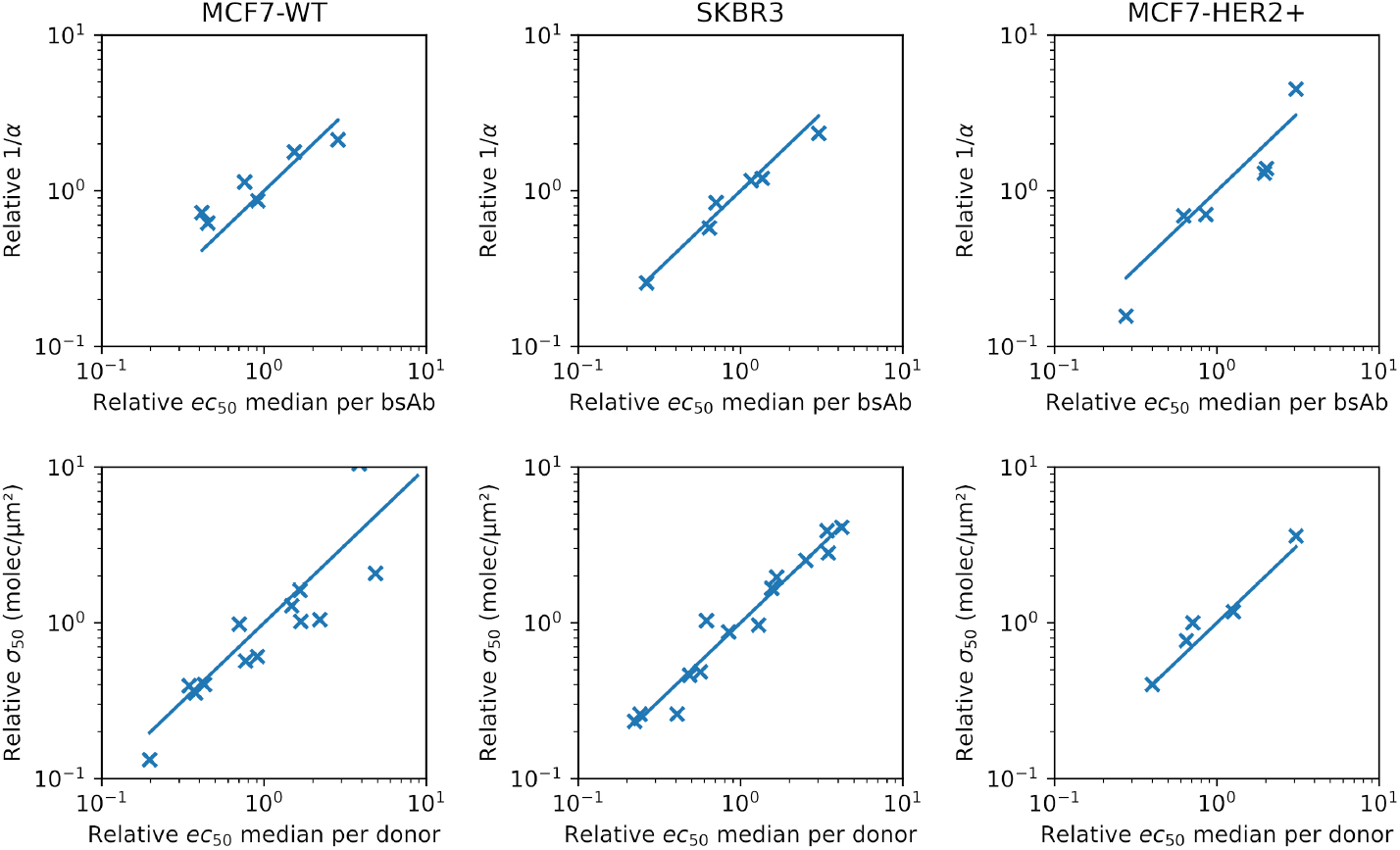
Comparison of relative fitting parameters and relative *ec*_50_ medians. Solid lines indicate *x* = *y*.

**Figure S16:**
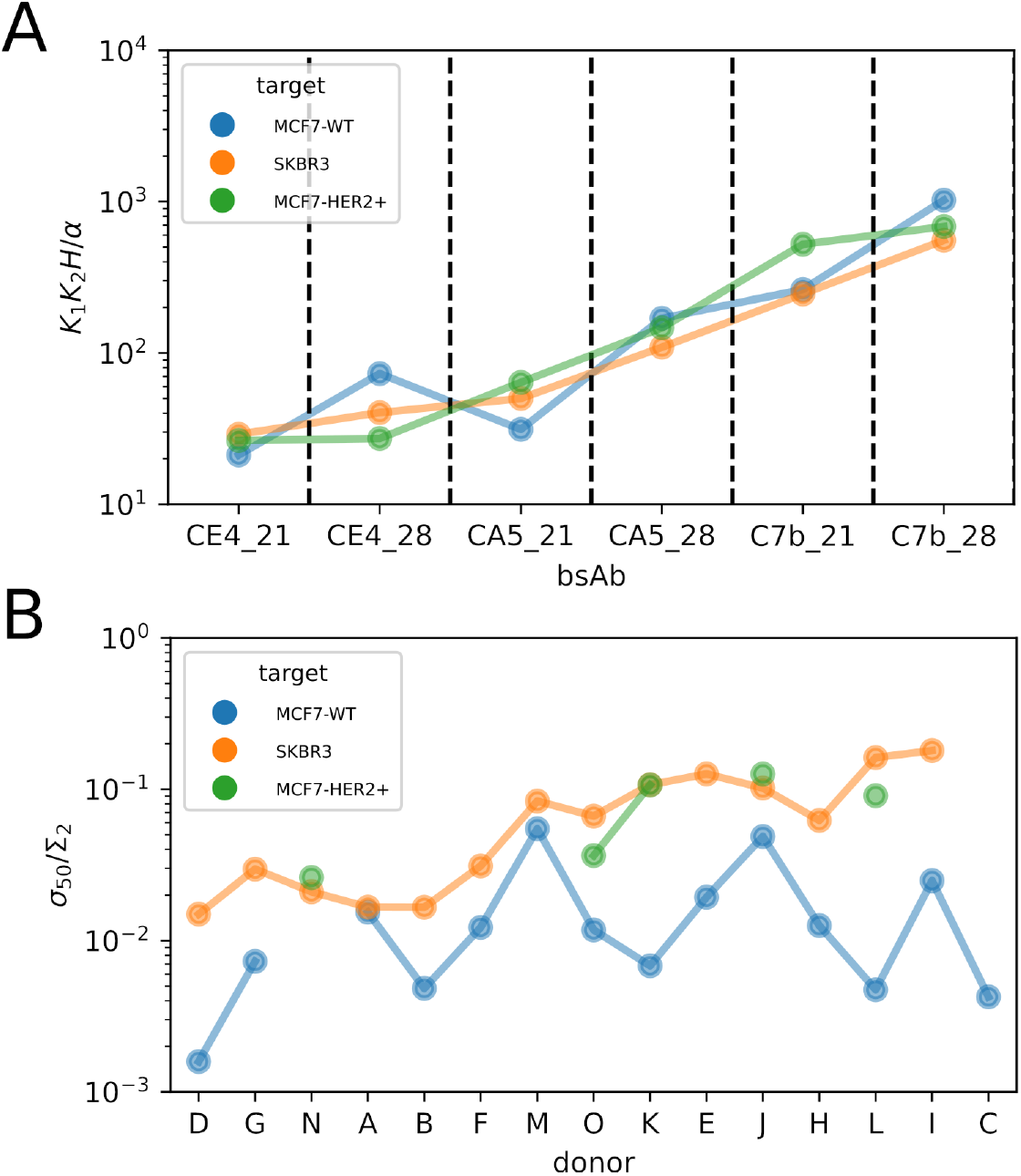
Estimated parameters for the dependence of EC_50_ on bsAbs or donors (see Eq. 4 of the Main Text). 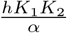 in *µ*m*×*(nmol/L)^2^ depends on the bsAb type. The dimensionless parameter *σ*_50_*/*Σ_2_ depends on the donor.

**Figure S17:**
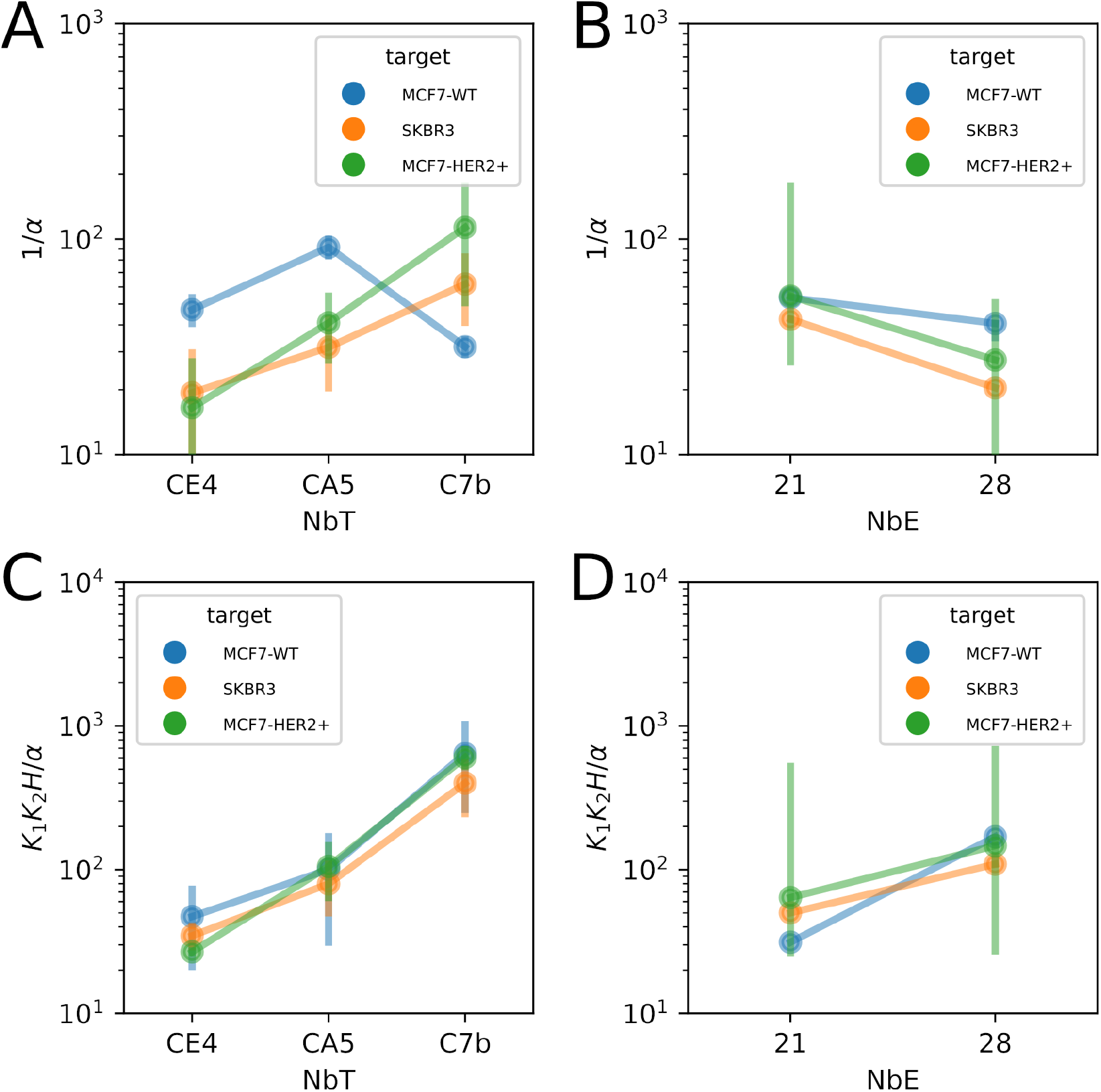
Estimated parameters for EC_50_ dependence on nanobody constituting the bsAb, on target or effector side. 1*/α* is dimensionless and 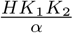 in *µ*m*×* (nmol/L)^2^. NbT: nanobodies against tumour cells. NbE: nanobodies against effector cells.

**Figure S18:**
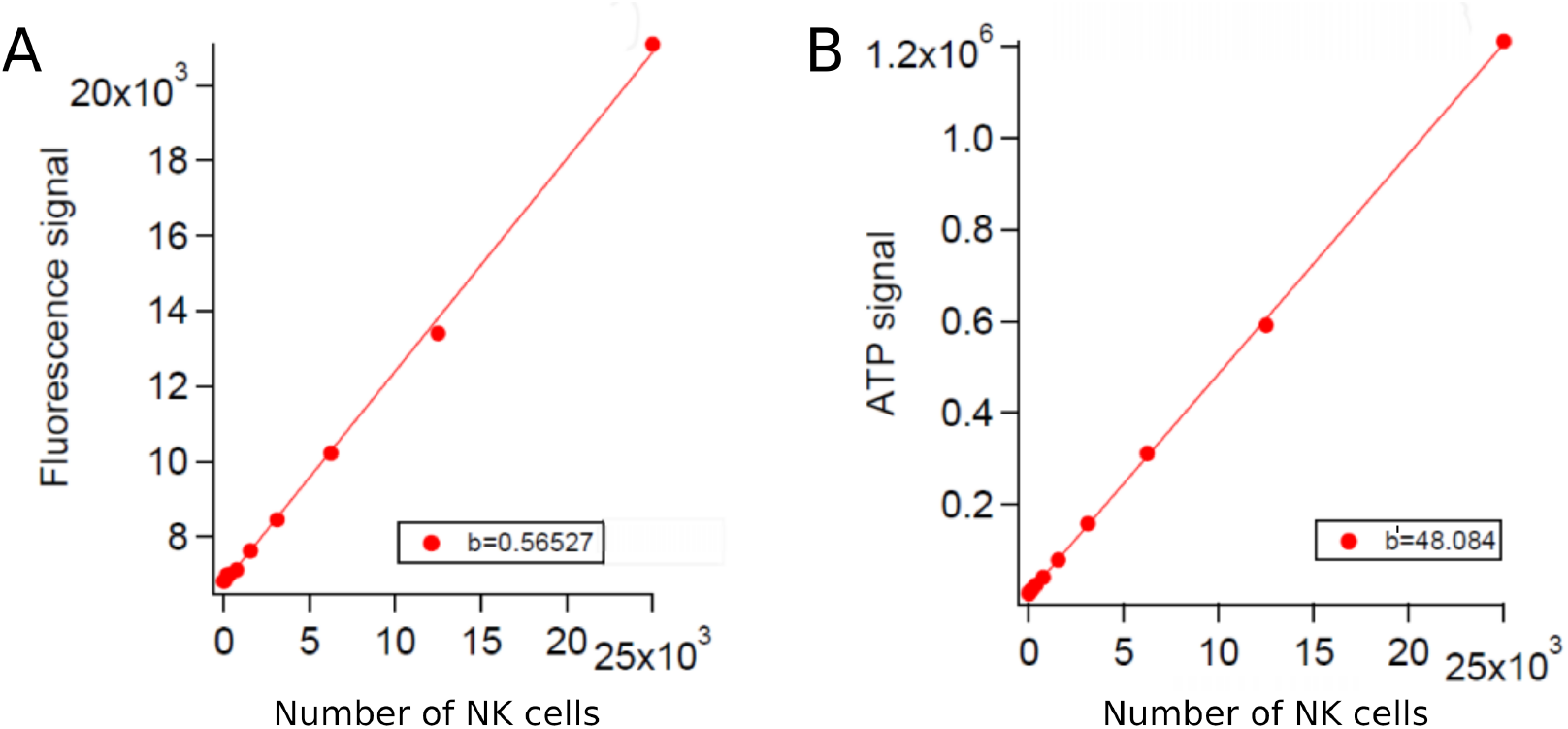
Calibration of NK cells luminescence for cell titer Glo assay. A. Relation between measured CFSE fluorescence signal and number of NK cells. B.Relation between luminescence ATP signal and number of NK cells. See Methods for details.

**Figure S19:**
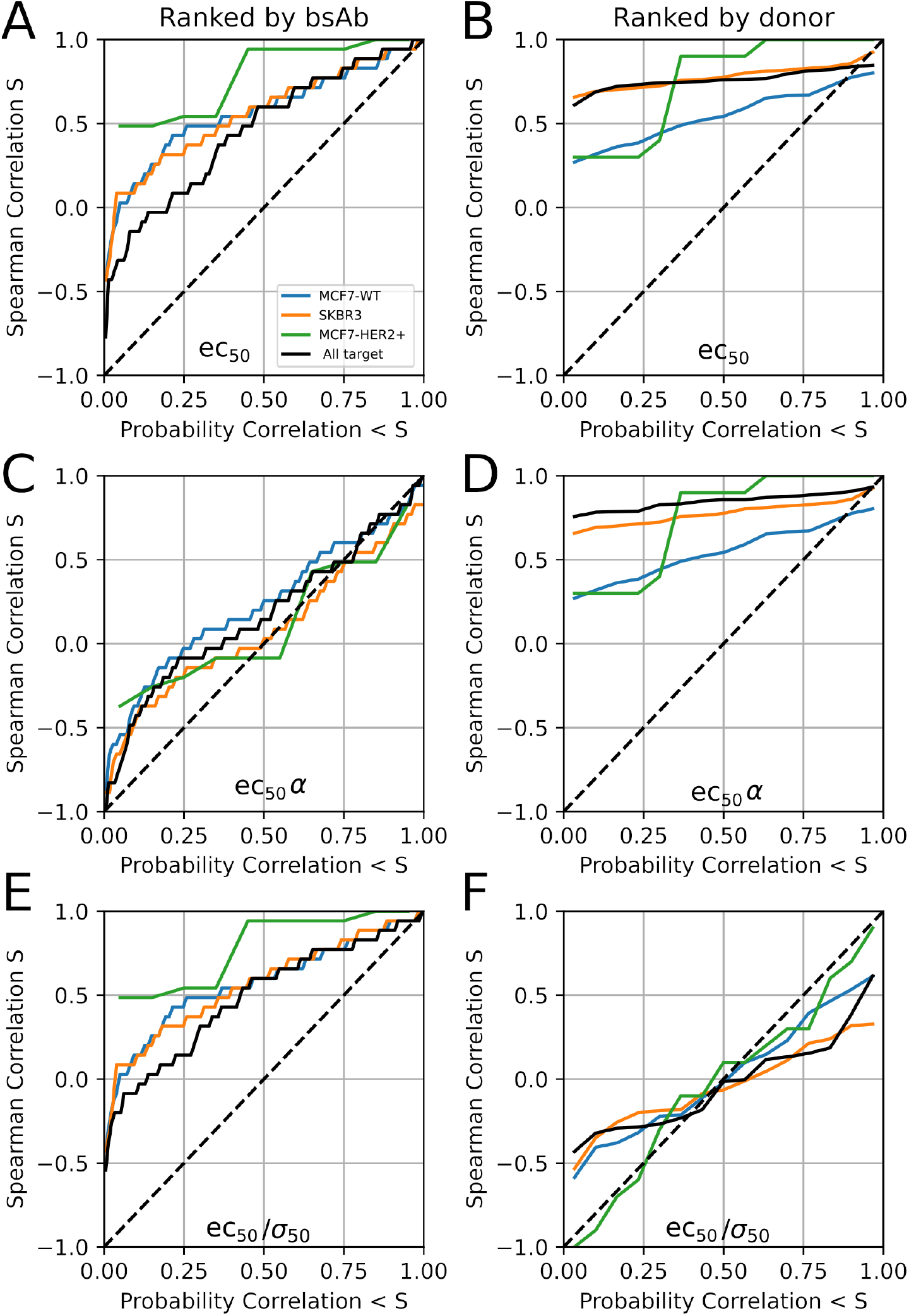
Spearman ranking based reduced potency *ec*_50_ – see definition in Tab. 1. (A,B), *ec*_50_*α* (C,D) and *ec*_50_*/σ*_50_ (E,F). A, C, E. ranking of bsAb for pairs of donors. B, D, F ranking of donors for pairs of bsAb. Dashed lines indicate a random order. See Tab. S4 for numbers.

**Figure S20:**
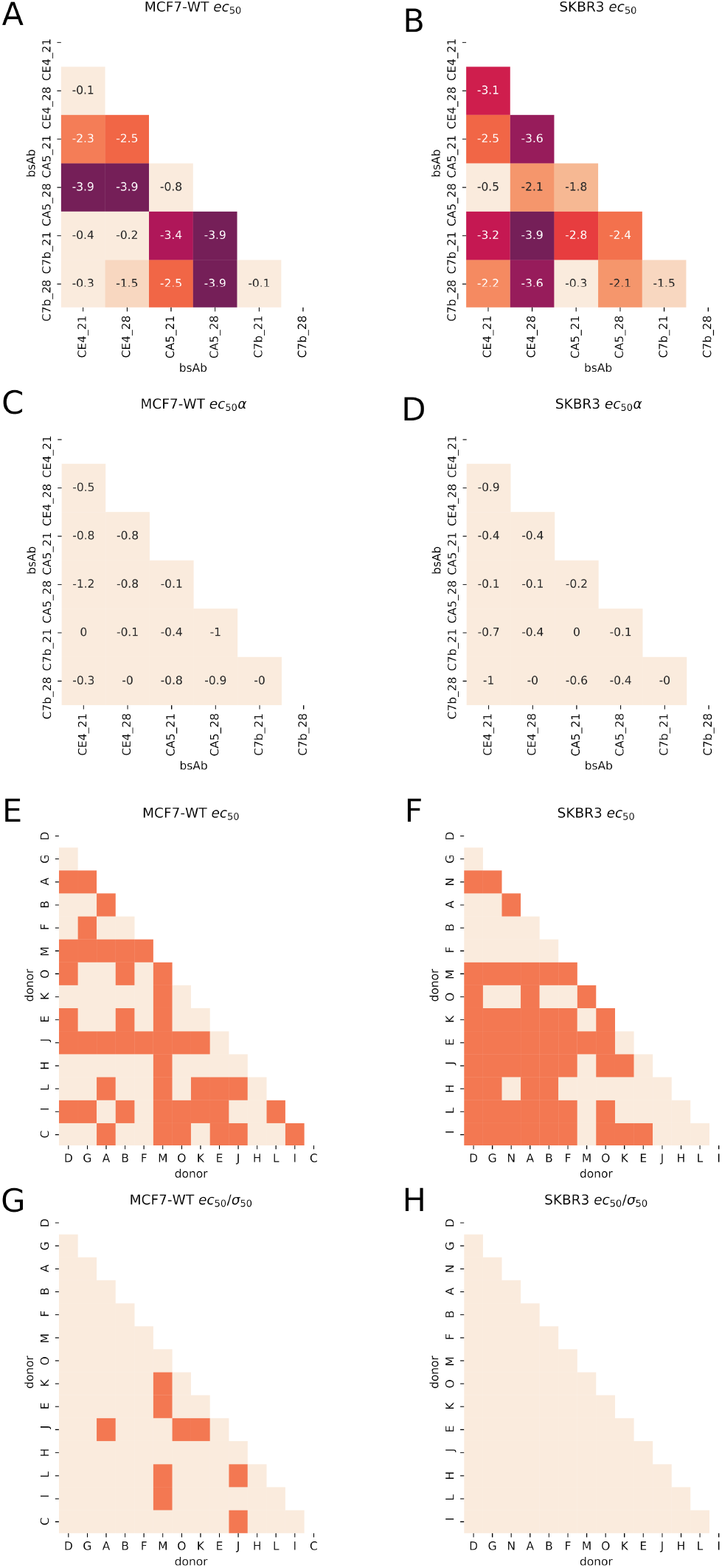
Wilcoxon pairwise comparison of reduced potency *ec*_50_ – see definition in Tab. 1. (A, B, E, F), *ec*_50_*α* (C, D) and *ec*_50_*/σ*_50_ (G, H). Left column: MCF7-WT. Right column: SKBR3. (A-D) ranked by bsAb. (E-H) ranked by donor.

**Figure S21:**
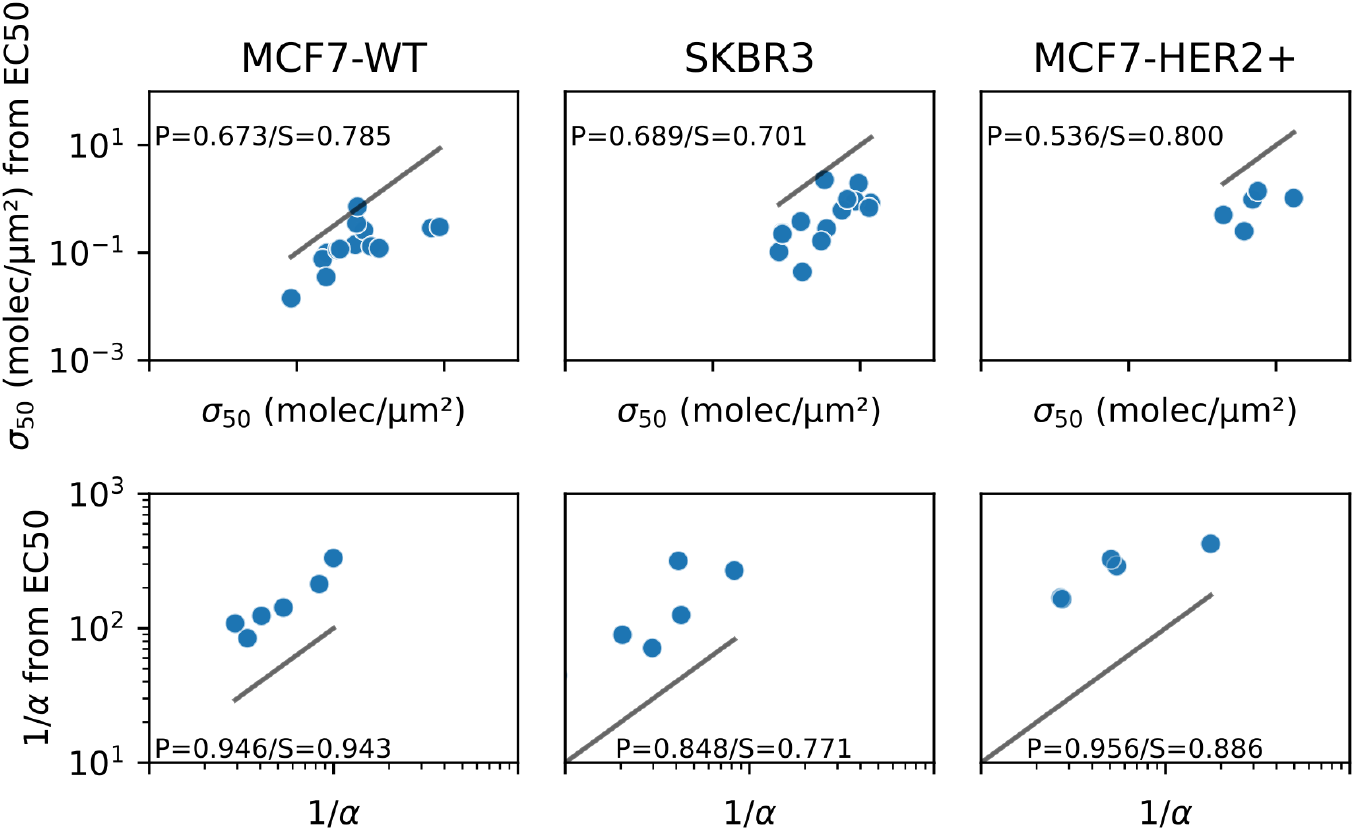
Correlation between best-fit values of parameters obtained from a global fit (horizontal axis) vs those from fits of EC_50_ (vertical axis). Solid lines indicate *x* = *y*.

**Figure S22:**
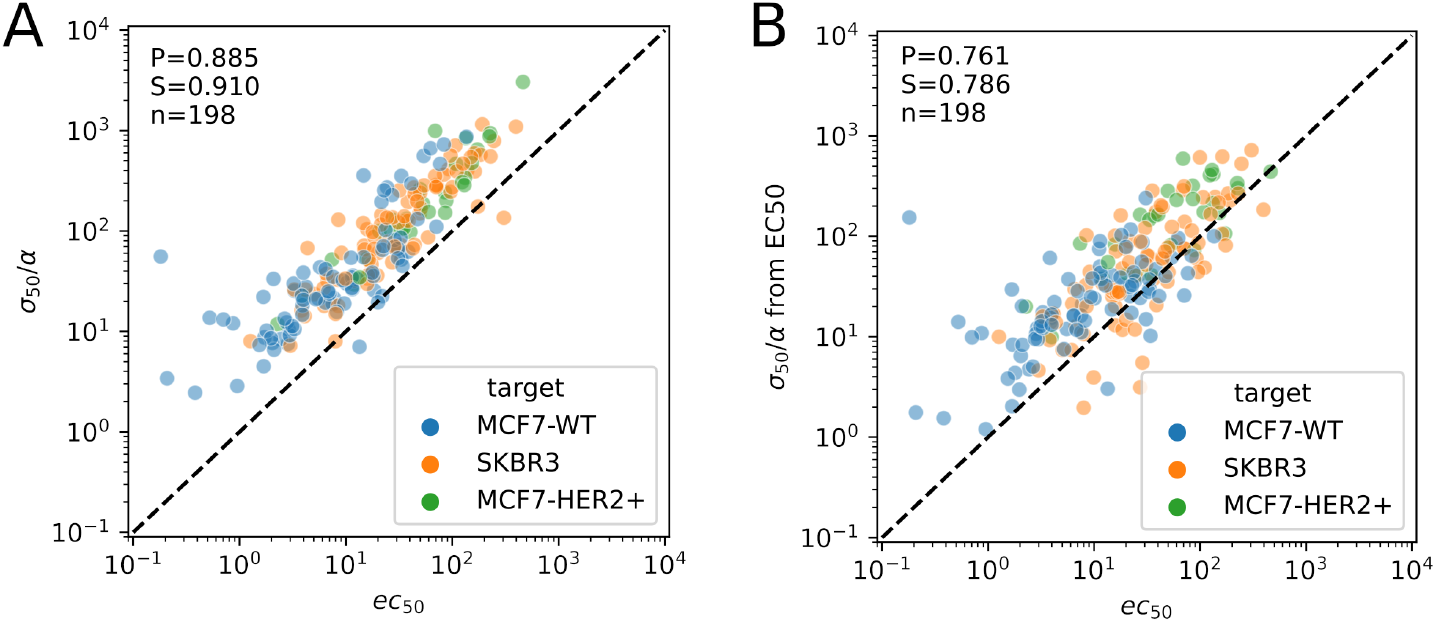
A. Relation between the rescaled potency *ec*_50_ (horizontal axis) and the quotient *σ*_50_*/α* (vertical axis). B. Relation between the quotient *σ*_50_*/α* obtained by the fitting of the full lysis data set (horizontal axis) or from fitting only EC_50_, vertical axis). Dashed lines represent *y* = *x*. Each point represents one Donor/bsAb/Target condition. P: Pearson coefficient. S: Spearman coefficient. n: number of points.

